# Optimized cDICE for efficient reconstitution of biological systems in giant unilamellar vesicles

**DOI:** 10.1101/2021.02.24.432456

**Authors:** Lori Van de Cauter, Federico Fanalista, Lennard van Buren, Nicola De Franceschi, Elisa Godino, Sharon Bouw, Christophe Danelon, Cees Dekker, Gijsje H. Koenderink, Kristina A. Ganzinger

## Abstract

Giant unilamellar vesicles (GUVs) are often used to mimic biological membranes in reconstitution experiments. They are also widely used in research on synthetic cells as they provide a mechanically responsive reaction compartment that allows for controlled exchange of reactants with the environment. However, while many methods exist to encapsulate functional biomolecules in GUVs, there is no one-size-fits-all solution and reliable GUV fabrication still remains a major experimental hurdle in the field. Here, we show that defect-free GUVs containing complex biochemical systems can be generated by optimizing a double-emulsion method for GUV formation called continuous droplet interface crossing encapsulation (cDICE). By tightly controlling environmental conditions and tuning the lipid-in-oil dispersion, we show that it is possible to significantly improve the reproducibility of high-quality GUV formation as well as the encapsulation efficiency. We demonstrate efficient encapsulation for a range of minimal systems including a minimal actin cytoskeleton, membrane-anchored DNA nanostructures, and a functional PURE (Protein synthesis Using Recombinant Elements) system. Our optimized cDICE method displays promising potential to become a standard method in biophysics and bottom-up synthetic biology.

## Introduction

Cellular life is enabled by countless interacting molecules and biochemical reactions with a high degree of interconnectivity and redundancy. Reconstituting cell biological processes using only their minimal functional units from the bottom-up is therefore greatly helpful to study cellular mechanisms on a molecular and mechanistic level.^1–3^ The field of bottom-up synthetic biology has gained a lot of traction over the last decade, an evolution synchronised with the emergence of several different consortia worldwide to lead the journey towards functional reconstitution of all basic cellular functions, culminating in the creation of a minimal synthetic cell.^4–7^

In this synthetic cell community, giant unilamellar vesicles (GUVs) are widely used as cell-sized, lipid bilayer-enclosed reaction compartments that can be visualized by real-time microscopy and directly manipulated using biophysical tools.^8–11^ Using GUVs as a basis for a functional synthetic cell requires encapsulation of different biological modules in a precise stoichiometry, consisting of a variety of biomolecules ranging in size and charge. However, state-of-the-art GUV fabrication methods are still far from ideal in establishing complex reconstituted systems. On the one hand, easy-to-implement and high-yield methods, such as natural swelling^12^, electroformation^13–16^, and gel-assisted swelling^17–20^, offer poor control over encapsulation efficiency and stoichiometry, and inconveniently contain the same solution on the in- and outside. On the other hand, emulsion-based techniques, in which GUVs are generated from water-in-oil droplets crossing an oil-water interface (using gravity, centrifugation, microfluidic devices or microfluidic jetting^21–27^), offer more control over GUV content and size monodispersity, but at the cost of being less reliable and more technologically advanced and therefore less accessible.

A promising method that is increasingly being used for complex reconstitutions is continuous droplet interface crossing encapsulation (cDICE). This double-emulsion based technique relies on the continuous transfer of capillary-generated water-in-oil droplets across an oil-water interface using centrifugal force.^28^ Requiring only easy-to-operate laboratory instrumentation, cDICE can in principle provide high yields while being less technologically demanding than microfluidic-based approaches and allowing for more control over size and encapsulated content than swelling methods.^28,29^ However, despite promising first outcomes, using cDICE for protein encapsulation has remained difficult, beyond a few specific cases^30–33^. At least in part, this is likely due to our lack in understanding of both the physical process of vesicle formation and of which parameters are essential to control tightly for the method to work robustly. Significant lab-to-lab variability and constant adaptations to the protocol devised by various labs^28–30,33^ have also made it hard to reproduce results across different institutions, leading to the technique being far from accessible.

Here, we aimed to gain a better understanding of the parameters influencing both vesicle formation and encapsulation efficiency in cDICE, allowing us to design an accessible, robust and reproducible workflow for different encapsulation needs. We show that control of environmental conditions is crucial for reliable formation of defect-free GUVs (*i.e.*, the vesicular membrane is uniform at optical length-scales and does not contain visible lipid pockets) at high yields. Furthermore, we demonstrate different approaches for enhancing the encapsulation efficiency of cDICE by changing the composition of the lipid-in-oil dispersion. We thus provide future users with a detailed protocol for GUV fabrication and a toolbox that can form a firm basis for further experiment-specific optimization. By reproducing key experiments across multiple labs in different locations and encapsulating a large variety of biological systems, from the encapsulation of purified proteins to the PURE *in vitro* transcription-translation system, membrane-anchored DNA origami, and bacteria, we show robustness and versatility of the method. Overall, we demonstrate that our improved cDICE protocol shows great promise for a wide range of complex reconstitution processes in the future, overcoming a major hurdle on the route towards functional minimal synthetic cells.

## Results

### Environmental control is essential for producing defect-free GUVs with cDICE

To improve the robustness of the cDICE method, we sought to systematically screen various experimental parameters that might influence GUV formation in cDICE. A typical cDICE set-up (Fig. 1a) consists of a rotating chamber containing two concentric fluid layers, an inner, lower-density lipid-containing oil phase and an outer, aqueous layer. The aqueous solution to be encapsulated is injected into the lipid-in-oil layer through a capillary, leading to the formation of water-in-oil droplets at the capillary orifice. As these droplets travel outward and traverse the interface of the oil with the outer aqueous phase, a bilayer is formed, yielding GUVs, collected in the outer layer of the system (Fig. 1a). GUV formation is thus dependent on the properties of all phases and on other experimental parameters, such as rotation speed and capillary size.^28^ When we sought to enhance the consistency of vesicle production in this inherently sensitive experimental system, the first striking improvement was made by using a chloroform-based lipid-in-oil dispersion^33^ as oil phase and preparing it in a humidity-free environment, *i.e.,* inside a glovebox. Without the use of a glovebox, GUVs were generated but the sample contained a lot of residual membrane material, such as free tubes and fluorescent aggregates, and the vast majority of GUVs showed visible fluorescent pockets or budding membrane structures (Fig. 1b). In contrast, when the lipid-in-oil dispersion was prepared in a glovebox, samples were much cleaner with most GUVs having quasi-spherical shapes without visible lipid pockets or budding membrane structures (Fig. 1c).

**Figure 1.**
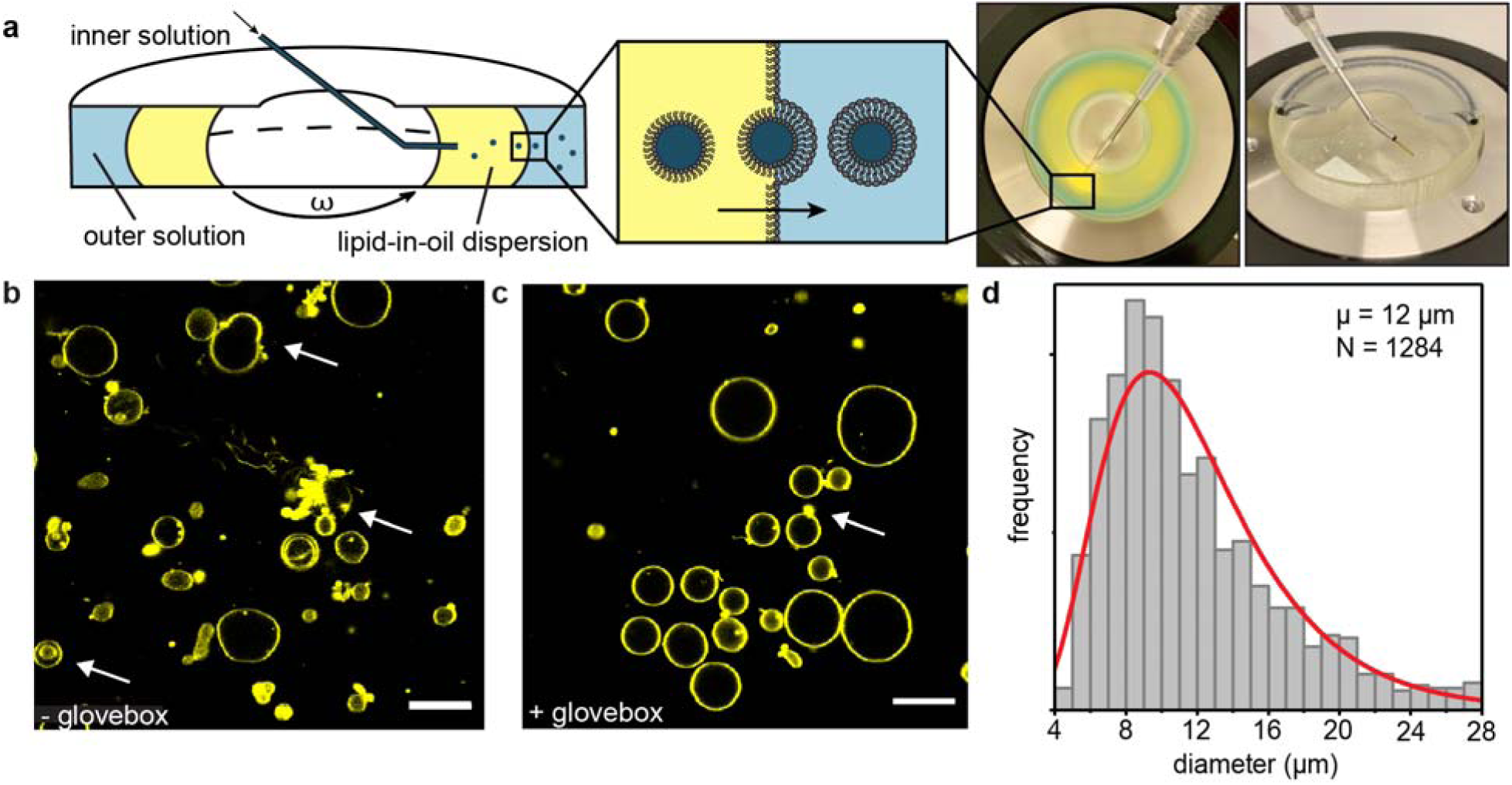
General overview of the cDICE technique and influence of environmental conditions. a. Cross-sectional schematic of the cDICE method. The centre image displays the 3D printed rotation chamber, with the different fluid layers coloured differently for illustration purposes. The rightmost image displays the custom-built spinning device that accommodates the 3D printed rotation chamber. The capillary is inserted using an adjustable magnetic base to allow spatial flexibility upon insertion. During experiments, this set-up is connected to a syringe and syringe pump. b. Representative field of view of GUVs formed using a chloroform-based lipid-in-oil dispersion prepared outside of the glovebox. ATTO 655 DOPE was used as a membrane stain and images were taken using confocal microscopy. Most GUVs contain artefacts in the lipid membrane, examples are indicated with arrows. Scale bar indicates 20 µm. c. Representative field of view of GUVs formed using the final protocol including the use of a glovebox. ATTO 655 DOPE was used as a membrane stain and images were taken using confocal microscopy. Most GUVs are spherical and possess a clean membrane and only a small population of GUVs still shows artefacts, as indicated with an arrow. Scale bar indicates 20 µm. d. Size distribution of GUVs made of DOPC lipids, obtained by the optimized protocol. The distribution is fitted to a log-normal function (red curve).

In line with this observation, preparation of the lipid-in-oil dispersion inside a glovebox also affected its macroscopic appearance: oil dispersions prepared in a humidity-free environment were transparent, while preparations outside a glovebox yielded visibly opaque dispersions, as quantified by turbidity measurements (A_350_ = 0.10 ± 0.05 vs 0.42 ± 0.10, SI Fig. 1). Furthermore, we analysed the lipid adsorption kinetics of the different oil dispersions using pendant drop measurements^34^, where a drop of aqueous solution is suspended in a lipid-in-oil mixture, mimicking the process happening at the orifice of the cDICE capillary. Without humidity control, interfacial tension decreased much faster (SI Fig. 2), indicating faster adsorption of lipids to the water-oil interface. In combination with the adverse effect on vesicle quality, our experiments suggest that presence of water in the lipid-in-oil dispersion interferes with vesicle formation and bilayer quality via changing the microscopic organization of the lipids and their adsorptive behaviour.

It is well known that humidity values change throughout the year, reaching highest values in summer. This seasonal dependency in daily relative humidity can be as large as several tens in percentage^35^, equivalent to the range of 40 - 75% that we observed in the lab. Given the importance of humidity in preparation of the lipid-in-oil dispersion, we extended environmental control to regulating humidity in the room where the cDICE experiments were performed by using a dehumidifier. Indeed, dehumidification down to 30 - 40% resulted in smaller variability between lipid adsorption kinetics as measured in pendant drop experiments (SI Fig. 2), indicating a more reproducible adsorption behaviour. In line with the lower variability found in lipid adsorption rates, dehumidification also proved to be essential for reliable production of clean vesicles throughout the year. Taken together, using a glovebox for preparation of the lipid-in-oil dispersion and storage of its components, and performing cDICE experiments in a continuously dehumidified room, resulted in a robust formation of clean GUVs.

In the original cDICE paper^28^, as well as in other follow-up studies^29,30,36,37^, injection capillaries were pulled from glass tubes to final orifice diameters of a maximum of 20 µm. Since we found these narrow glass capillaries to be a significant source of experimental variation and problems due to easy clogging of the orifice, we explored if using commercially available fused silica capillary tubing with larger diameters (25, 50 and 100 µm) would allow for even more consistent results, as previously used by Litschel *et al.*^33^. We found that using all three capillary sizes, our chloroform-based lipid-in-oil dispersion and optimized workflow led to high yields of GUVs with a mean diameter of 12 µm and coefficient of variation of 47% (Fig. 1d). The size distributions of the GUVs did not significantly change across the different capillary sizes (SI Fig. 3) and they were broader than the ones previously obtained for smaller orifice sizes^28^. However, the lack of control over GUV size is compensated by a much-improved reliability of encapsulation and GUV formation due to avoidance of clogging, in particular for 100 µm fused silica capillaries. Other capillary materials were also successfully used, *i.e.*, 100 µm PEEK capillary tubing (Fig. 4a *i* and Fig. 4b). Within the margins of rotation speed tested (1000 – 2900 rpm), no precise control of speed is necessary in order to get robust GUV formation for all orifice diameters, with size distributions in an ideal range for bottom-up reconstitution of eukaryotic cells (SI Fig. 3). In terms of yield, the absolute number of GUVs obtained using the optimized cDICE protocol is generally variable, but from the average number of GUVs visible per field of view, we estimate the absolute number of GUVs to reach well over 1000 vesicles in most experiments.

### Unilamellarity of cDICE-produced GUVs

Many reconstitution experiments require unilamellar lipid membranes, as this determines permeability and mechanical properties of the GUV and is needed for insertion of transmembrane proteins, including pore proteins, into the bilayer. Therefore, we next aimed to investigate if our GUV membranes were unilamellar by monitoring insertion of alpha-hemolysin, a protein that assembles a heptameric pore structure in the lipid membranes with a diameter of 14 Å, through which small molecules can pass and which is highly sensitive to the thickness of lipid bilayers.^38,39^ As a tracer, we encapsulated 5 µM of the fluorescent dye Alexa Fluor 488 (643 Da) and we immobilized the GUVs to aid long-term imaging by adding a polyisocyanide hydrogel^40^, which transitions from a liquid to a gel phase when warmed up from 4 °C to room temperature. After that, alpha-hemolysin was added to the chamber and fluorescent imaging was immediately started. Within minutes following alpha-hemolysin addition, all GUVs observed started to lose their fluorescent content and all had lost 50% of their content after ∼ 20 minutes (Fig. 2a, top row; Fig. 2b, red curve). In stark contrast, when only alpha-hemolysin buffer was added to the GUVs as a control, fluorescent molecules were clearly retained within all GUVs (Fig. 2a, bottom row; Fig. 2b, blue curve). This indicated that loss of GUV content was due to pore formation and hence membrane unilamellarity. Furthermore, individual GUV membrane intensities normalised by the population’s mean membrane intensity are consistently distributed around unity, indicating a homogenous lamellarity over the GUV population (Fig. 2c). Taken together, our results clearly show that the cDICE method produces unilamellar GUVs.

**Figure 2.**
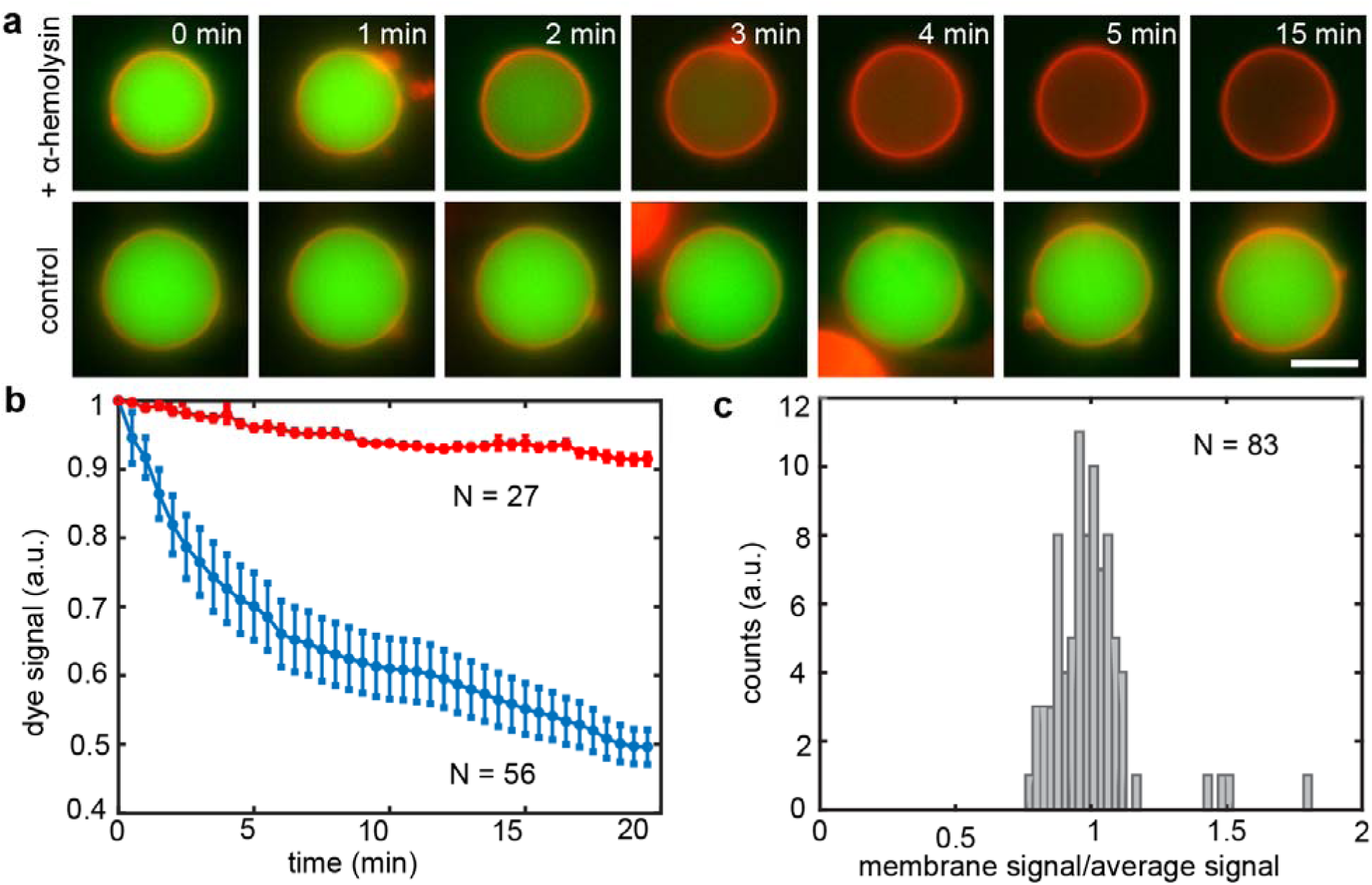
Incorporation of alpha-hemolysin pore protein demonstrates unilamellarity of GUV membrane. a. Fluorescence microscopy images of single GUVs prepared using a chloroform-based lipid-in-oil dispersion showing different membrane permeability in presence (top row) or absence (bottom row) of alpha-hemolysin. When the pore protein is added to the lipid membrane (red, rhodamine-PE fluorescence signal), the encapsulated fluorescent dye (green, Alexa Fluor 488) is released in the outer environment within a few minutes. When only alpha-hemolysin buffer is added as a control instead, fluorescent molecules are retained within the GUV volume. Scale bar indicates 5 µm. b. Quantitative analysis of GUV fluorescent content loss over time. In presence of alpha-hemolysin (blue curve), Alexa Fluor 488 signal intensity decreases down to 50% of the initial value within the first 20 minutes, while in absence of pores (red curve) only a minor decrease (< 10%), likely due to photobleaching, is detected. c. Histogram showing GUV membrane fluorescence intensities compared to the overall GUV population.

### Improvement of encapsulation efficiency

To allow for complex reconstitution experiments, it is essential to have control over the encapsulation of functional biomolecules in the right stoichiometric ratios. We probed the encapsulation efficiency of our improved cDICE protocol by encapsulation of the cytoskeletal protein actin, a broadly used protein in the synthetic biology field.^41^ While all experiments using our optimized cDICE protocol resulted in successful encapsulation of monomeric actin in GUVs at high vesicle yields, automated analysis of actin fluorescence at the equatorial plane of the GUV from confocal fluorescence imaging surprisingly revealed a substantial fraction of GUVs with very low actin content, indicating that many of the formed vesicles were seemingly empty (23%, Fig. 3a, SI Fig. 4). We tested if the encapsulation efficiency could be improved by using different lipid-in-oil mixtures. We reasoned that the encapsulation efficiency may depend on the lipid adsorption kinetics, as it has been reported earlier that the dispersion methods of lipids had a strong effect on their adsorptive behaviour.^42^ Therefore, we investigated the effect of lipid dispersion strategy on adsorption kinetics (by pendant drop measurements) and on GUV formation and encapsulation efficiency for three lipid mixtures: lipids dispersed as aggregates in a 80:20 mixture of silicon and mineral oil using chloroform as mentioned above, a similar dispersion of lipid aggregates but using decane instead of chloroform, and a lipid-chloroform solution in mineral oil only. Chloroform and decane serve as good solvents for the lipids, while the lipids do not dissolve in the oil. This way, we aimed to produce different lipid-in-oil dispersions with various aggregation states, with the mineral oil dispersion having smallest aggregate size, and both chloroform- and decane-based lipid dispersions having larger aggregate sizes.^42^

**Figure 3.**
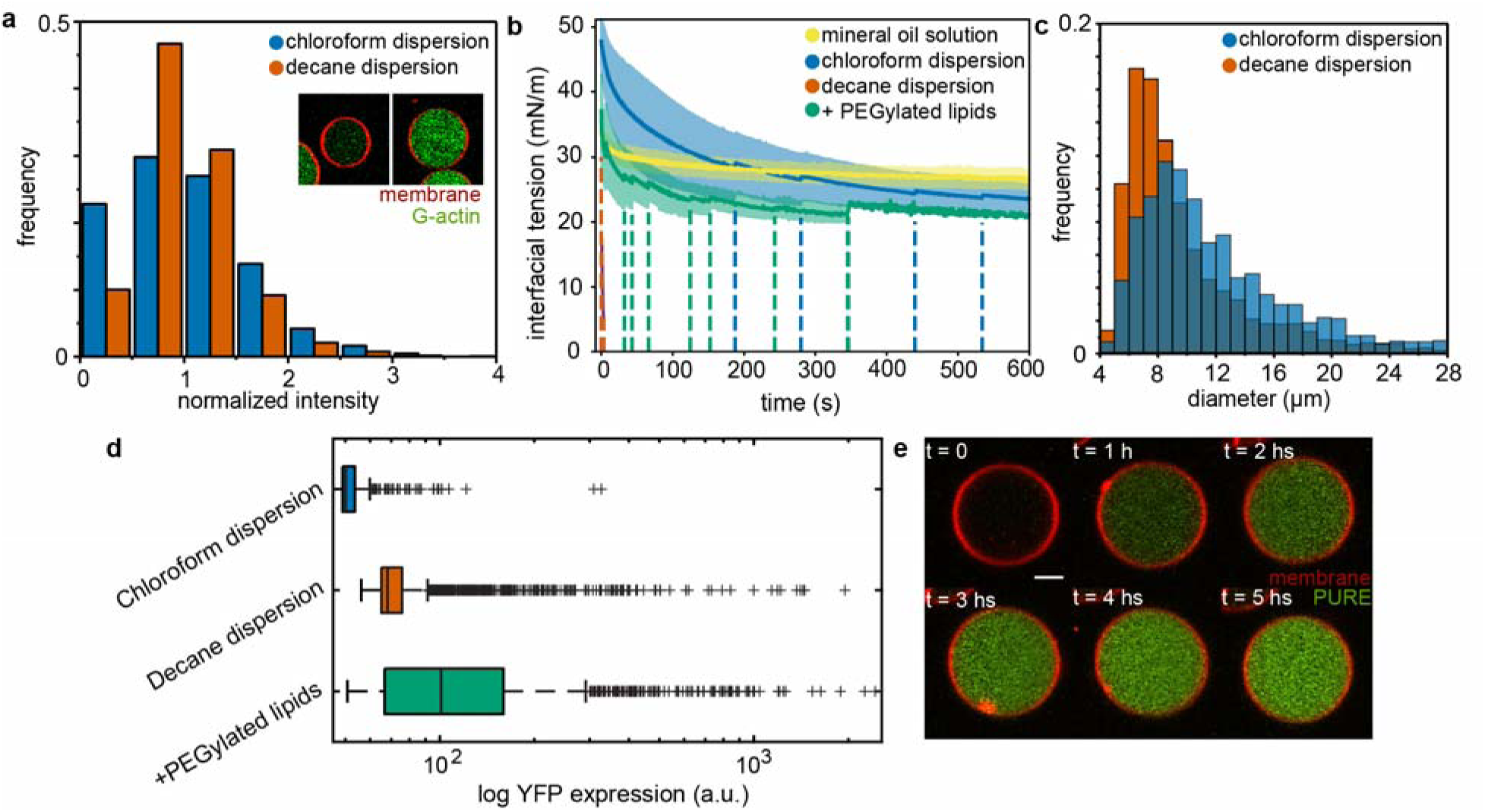
Improved encapsulation by tuning of the lipid-in-oil dispersion. a. Encapsulation efficiency of G-actin using a chloroform-based lipid dispersion (blue) and decane-based lipid dispersion (green). The first bin represents GUVs with very low fluorescence intensity, and represents 23% of the population for the chloroform-based lipid dispersion and only 10% for the decane-based lipid dispersion. b. Interfacial tension decrease measured for a pendant droplet of G-buffer in different lipid-in-oil mixtures. Solid lines show averaged data with standard deviation for a lipid-chloroform solution in mineral oil only (orange, n = 9), dispersed lipid aggregates using chloroform (blue, n = 13) or decane (green, n = 7) in silicone oil:mineral oil 80:20 and a chloroform-based lipid-in-oil dispersion with 0.01 mol% of PEGylated lipids (yellow, n = 9). The dashed lines indicate individual events where the droplet fell off, which gave rise to apparent jumps in the averaged curves. c. Size distribution of GUVs made using a chloroform-based lipid dispersion (blue) and decane-based lipid dispersion (green). d. Box plots of the YFP expression after five hours of incubation in GUVs obtained using dispersed lipid aggregates using chloroform (blue), decane (green), and a chloroform-based lipid-in-oil dispersion with 0.01 mol% of PEGylated lipids (yellow). The boxes represent IQR (25th–75th percentiles), the centre line indicates the median and the whiskers extend to the maximum and minimum value excluding outliers. Outliers are individually indicated using plus symbols. e. Time-lapse images of YFP expression in a single GUV using a chloroform-based lipid-in-oil dispersion with 0.01 mol% of PEGylated lipids. Scale bar indicates 5 µm.

First, we confirmed the aggregation state of the lipids by absorbance measurements. Indeed, the mineral oil dispersion was much less turbid (A_350_ = 0.03 ± 0.01) than the chloroform- or decane-based dispersion (A_350_ = 0.10 ± 0.05 and A_350_ = 0.20 ± 0.12 respectively, SI Fig. 2), indicating the presence of larger aggregates in the latter two. Pendant drop measurements showed that dispersing lipids as aggregates using chloroform resulted in fast lipid adsorption (Fig. 3b, blue curve), indicating fast monolayer formation. The decane-based lipid dispersion resulted in even faster adsorption, with all droplets detaching within several seconds (Fig. 3b, green curve). In contrast, lipids dispersed in mineral oil exhibited a slower and smaller decrease of interfacial tension (Fig. 3b, orange curve), meaning slow adsorption of lipids to the oil-water interface and a small coverage of the final interface. In line with the idea that faster stabilization of the oil-water interface by faster lipid adsorption leads to more robust monolayer formation, we observed no GUV formation when using lipids dispersed in mineral oil, whereas experiments using lipids dispersed as aggregates in a 80:20 mixture of silicon and mineral oil using chloroform or decane gave large GUV yields (SI Fig. 4).

We then tested if the fast-adsorbing decane mixture could improve the encapsulation efficiency of cDICE. In stark contrast to the encapsulation of G-actin using chloroform as an organic solvent, using a decane-based lipid dispersion resulted in a significant decrease of the fraction of seemingly empty vesicles (10% versus 23%, Fig. 3a, SI Fig. 4). Although large differences in both adsorption kinetics and encapsulation efficiency can be observed between decane- and chloroform-based lipid-in-oil dispersions, they yield GUVs similar in size distribution, size polydispersity, and visual membrane cleanliness (Fig. 3c, SI Fig. 4). We also note that the lipid adsorption behaviour of the chloroform-based dispersion is highly variable, much more so than for decane-based lipid dispersions or lipids dispersed in mineral oil only (Fig. 3b). Since the lipid dispersions are metastable mixtures and chloroform readily evaporates under ambient conditions, changes to their composition happen on timescales similar to the experimental runtime. Indeed, time-dependent absorbance measurements indicated a rapid change in turbidity on the time scale of minutes, confirming this intrinsic instability of chloroform-based lipid dispersions (SI Fig. 5).

Efficient encapsulation is particularly important for reconstitution of cell-free gene expression reactions (*in vitro* transcription-translation systems) within GUVs, as the relative stoichiometry of their components has to be rather closely retained for optimal functioning.^43^ Functionality might further be affected by possible hydrophobic interactions of the protein components with organic solvents during encapsulation, although some groups already successfully encapsulated *in vitro* transcription-translation systems with emulsion-droplet transfer-^44–46^ and microfluidic-based methods^47^. To our knowledge, functional encapsulation of a cell-free gene expression (*e.g.*, the Protein synthesis Using Recombinant Elements (PURE) system^48^) has never been demonstrated for GUVs produced with the cDICE method. We therefore explored if we could encapsulate the PURE system using our improved cDICE protocol. To this end, GUVs encapsulating PURE*frex*2.0, a commercially available PURE system, along with a linear DNA construct coding for yellow fluorescent protein (YFP), were produced using both a chloroform-based lipid dispersion and a decane-based lipid dispersion. Gene expression in GUVs incubated at 37 °C was monitored by imaging YFP production within the GUV lumen over time. We observed that the different dispersion strategies used for GUV fabrication influenced the level of gene expression: the distribution of luminal fluorescence intensity after five hours of gene expression employing decane-based lipid aggregates showed improved gene expression levels compared to the encapsulation using chloroform-based lipid aggregates, which barely yielded any expressing GUVs at all. Nevertheless, both levels and numbers of expressing GUVs were still very low (Fig. 3d and SI. Fig. 6).

In addition to the lipid dispersion strategy, the lipid composition of the bilayer membrane can also alter adsorption kinetics and hence improve encapsulation efficiency. In particular, PEGylated lipids, lipids with a flexible poly(ethylene) glycol (PEG) linker, are often proposed to boost robust vesicle formation for various protocols.^17,49–51^ We therefore investigated if doping the vesicular membrane with 0.01 mol% 18:0 PEG2000 PE could improve encapsulation of the PURE system when using cDICE. The presence of PEGylated lipids slightly increased the adsorption rate of lipids to the oil-water interface (Fig. 3b, yellow curve). Interestingly, doping the membrane with 0.01 mol% PEGylated lipids greatly enhanced expression of the encapsulated PURE system and resulted in the highest expression levels and in a large population of GUVs expressing YFP (Fig. 3d, e and SI. Fig. 6). These results show that optimization of encapsulation efficiency both via lipid dispersion and lipid composition is crucial to allow for functional reconstitution of complex reactions such as the PURE system in GUVs made using cDICE.

### Proof-of-concept experiments illustrate versatility of the optimized workflow

Finally, to investigate the broad applicability of our improved cDICE method, we aimed to reconstitute a wide range of minimal systems inside cDICE-made GUVs (Fig. 4). First, we encapsulated a minimal, branched actin network. In eukaryotic cells, the actin cortex is the protein machinery responsible for cell division.^52,53^ Reconstitution of a functional actin cortex anchored to the inner leaflet of the GUV membrane therefore offers an attractive route to induce GUV constriction, and possibly membrane fission, in synthetic cells. Our minimal actin cortex consisted of actin together with the verpolin homology, cofilin, and acidic domain of the Wiscot-Aldrin Syndrome protein (VCA), the Arp2/3 complex, an actin nucleator responsible for promoting formation of a branched actin network at the cell membrane^54,55^, and profilin. VCA was His-tagged to be able to bind to DGS-NTA(Ni) lipids in the membrane.^56,57^ Profilin was used to prevent actin polymerization in the GUV lumen and promote localized nucleation at the membrane.^58,59^ As a result, actin displayed a clear localization at the GUV membrane (Fig. 4a *i*, SI Fig. 7a), similarly to what was obtained using other GUV fabrication methods.^60,61^ In absence of membrane anchors and nucleators, actin was uniformly distributed within the GUV volume (Fig. 4a *i*, SI Fig. 7a).

**Figure 4.**
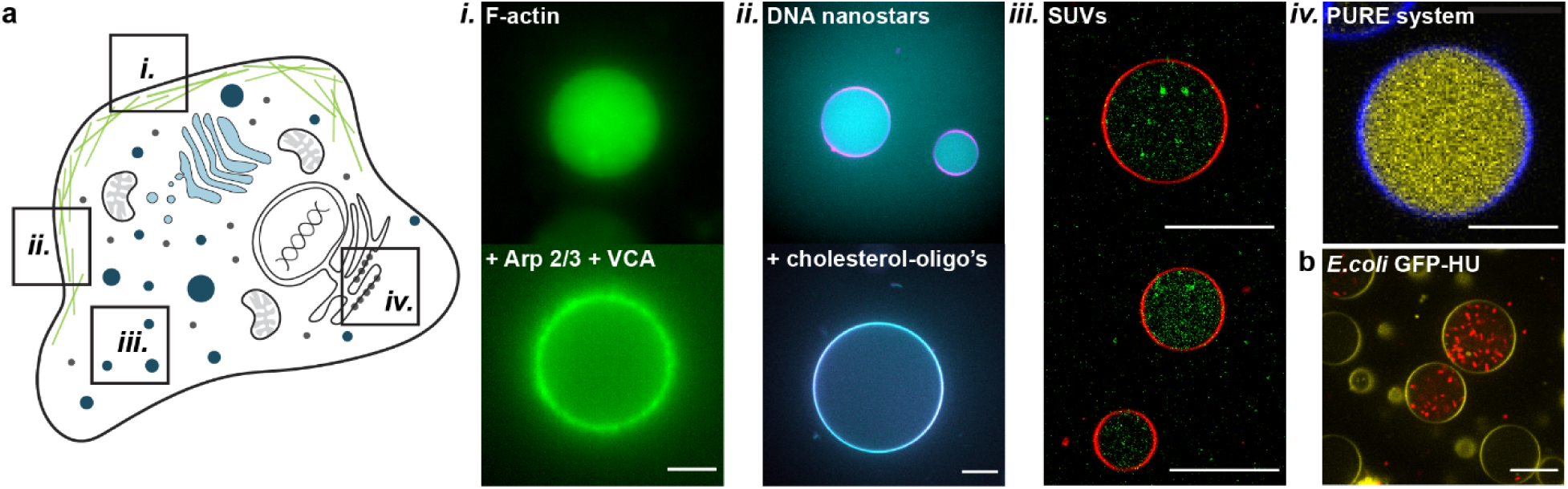
Proof-of-concept experiments showing versatility of cDICE and its applicability for the synthetic cell community. a. Overview - GUVs as artificial membrane systems to mimic cellular membranes and membrane interactions.

i. Reconstitution of a minimal actin cortex inside a GUV, nucleated at the vesicular membrane by the Arp2/3 complex, the C-terminal VCA domain of WASp, and profilin. Scale bar indicates 5 µm.
ii. Encapsulation of DNA origami nanostructures, freely diffusing inside the GUV lumen and capable of membrane localization upon addition of 2 µM of cholesterol-oligonucleotides. Scale bar indicates 15 µm.
iii. Encapsulation of SUVs inside GUVs to form a multicompartmentalized system. Scale bars indicate 20 µm.
iv. Encapsulation of PURE*frex*2.0 and DNA encoding for YFP. Scale bar indicates 10 µm. b. Encapsulation of GFP-HU expressing *E. coli* bacteria. A large number of bacteria could be observed inside the GUV lumen, clearly viable as evident from their motility. Scale bar indicates 20 µm.

As a synthetic mimic of the cellular actin cortex, we encapsulated DNA origami nanostructures^62^ that are capable of lateral polymerization. These four-armed DNA assemblies (SI Fig. 8) diffuse freely in the lumen of the GUV but were efficiently recruited to the membrane upon co-encapsulation of a cholesterol-oligonucleotide membrane anchor that binds single-stranded DNA sites on the origami (Fig. 4a *ii*, SI Fig. 7b). We also successfully encapsulated small unilamellar vesicles (SUVs, ∼ 100 nm diameter)^9^, mimicking multicompartmental cellular systems (Fig. 4a *iii*, SI Fig. 7c). In the future, these compartments could be designed to trigger or sustain intravesicular reactions, allowing control over biochemical reactions inside the GUV lumen.^63–65^

Furthermore, as mentioned above, our cDICE method can be used to encapsulate a functional *in vitro* transcription-translation system (the PURE system), provided PEGylated lipids are included in the lipid mixture (Fig. 4a *iv*). The broad applicability of cDICE is further demonstrated by the successful encapsulation of objects that are large compared to the GUV size, *i.e.* entire *E. coli* bacteria (Fig. 4b, SI Fig. 7d). Cylindrical in shape, with a length of approximately 3 µm and a diameter of 1 µm^66^, these are several orders of magnitude larger than even many DNA origami structures. The bacteria were clearly mobile inside the GUVs (SI Mov. 1), showing that the cDICE process does not significantly affect their viability. Encapsulating live bacteria inside synthetic cells could be a promising route to combine ‘the best of both worlds’, *e.g.*, photosynthetic cyanobacteria could be repurposed as ‘chloroplasts’ for the synthetic cell, similar to a recent study which included chloroplasts isolated from plant cells^67^.

Overall, the improved cDICE method is shown to be capable of encapsulating a variety of functional minimal systems related to cell mechanics, cell metabolism and gene expression, all required for the generation of a synthetic cell.

## Discussion

A good understanding of the parameters influencing the GUV formation process in cDICE is crucial, especially for design of reconstitution experiments beyond first proof-of-concept experiments. Here, we showed that tight control over the lipid-in-oil mixture is key to successful and reproducible GUV formation. We found that membrane quality, which affects mechanical measurements and quantitative fluorescence analysis, was strongly improved by environmental control over preparation and handling of the lipid-in-oil dispersion, notably handling the lipid dispersion in a humidity-free environment (*i.e.*, a glovebox) and decreasing humidity to 30% during vesicle formation. We hypothesize that air humidity affects bilayer formation via changing the microscopic aggregation state of the lipid-in-oil mixture, and thereby the lipid adsorption behaviour.^68^ Importantly, we also demonstrated the unilamellarity of the formed GUVs by correct insertion of alpha-hemolysin to allow pore formation. Although the appearance of the GUV membranes was visibly improved upon environmental control, a common concern remains the possible presence of residual oil traces in the membrane, which could alter membrane characteristics. In line with visual inspection and our alpha-hemolysin experiments, and as observed in previous work, cDICE-formed GUVs are unlikely to have large traces of oil persisting in the membrane.^28,29^ Interestingly, more recent work indicates that the possible presence of residual oil in vesicular membranes does not significantly alter their membrane properties compared to electroformed GUVs.^69–71^ Altogether, this makes vesicle formation with the improved protocol compatible with reconstitution experiments requiring clean unilamellar membranes, such as studies involving membrane mechanics, insertion of pore proteins or membrane permeability.

Furthermore, we showed that the dispersion state of the lipids is crucial for efficient GUV formation using cDICE. As other existing protocols show, many different lipid-in-oil mixtures can be used for GUV formation.^28–30,33,42^ In particular, Claudet *et al.*^42^ found lipids dispersed as aggregates in an oil phase to promote more efficient bilayer formation. We provide experimental evidence that indeed the lipid bulk aggregated state strongly influences adsorption kinetics and thereby vesicle formation, supporting and explaining the observations of Claudet *et al.*^42^. Our tensiometry findings also indicate that not solely adsorption speed is of importance for proper bilayer formation, but the structure and content of the lipid aggregates is just as crucial for mono- and bilayer formation. Hence, having lipids dispersed as aggregates alongside humidity control is essential for clean GUV formation. This indicates a non-trivial relation between lipid properties, lipid dispersion state, adsorption kinetics and the final membrane quality. Adsorption speed as measured by pendant drop experiments can therefore not be used as a stand-alone quantity to assess whether a given lipid-in-oil mixture will support GUV formation in cDICE.

By tuning the lipid-in-oil dispersion with different organic solvents or different types of lipids, the encapsulation efficiency of cDICE could be improved. Faster lipid adsorption when using a decane-based dispersion as compared to using a chloroform-based dispersion, led to a better G-actin encapsulation. For functional encapsulation of the PURE system on the other hand, the presence of PEGylated lipids proved to be crucial. This cell-free expression system has a complex molecular composition, and all the individual components need to be present in order to yield a functional readout. While addition of PEGylated lipids has proven to be very effective for encapsulation of the PURE system with cDICE, it should be noted that PEGylated lipids can have adverse effects on protein functionality and membrane physicochemical behaviour, as the polymer chains introduce crowding and steric repulsion of components from the membrane as well as affect the membrane thickness.^72^ In this case, our experiment suggest that depending on the encapsulated species, PEGylated lipids can be avoided and high encapsulation efficiencies can be reached instead by changing the solvent.

Our cDICE protocol robustly yields GUVs with an average diameter of 12 µm and coefficient of variation of 47%. This size distribution was robust to changes in rotation speed and capillary diameters from 25 - 100 um. This consistency over differences in these two central parameters implies that the workflow we have adopted lies in the jetting regime.^73^ A jet at the capillary orifice is broken up into a polydisperse droplet population due to the Rayleigh instability in combination with the centrifugal force applied in cDICE.^73^ A high degree of polydispersity can be advantageous for bulk assays to screen multiple conditions in one single experiment^74,75^, but undesirable for other applications. As Abkarian *et al.*^28^ showed, decreasing the capillary diameter to values around 10 µm or using an additional inner fluid layer to decrease shear forces are viable strategies to achieve more precise size control. However, using these small orifice sizes poses other problems, including fast clogging of small diameter capillaries, rendering the method much less reliable. Here, we demonstrate that to reproducibly encapsulate viscous solutions containing a high concentration of polymerizing protein, as when encapsulating concentrated actin solutions, it is advantageous to use a larger capillary.

Taken together, the optimized workflow laid out in this research, focusing mainly on control of humidity and design of the lipid-in-oil dispersion, enables the generation of bespoke GUVs at good yields and with high encapsulation efficiency. We showed that encapsulation was compatible with molecular membrane anchors such as the cholesterol-oligonucleotide anchors used with DNA origami and a minimal actin cortex, while maintaining functionality even for complex systems like the PURE system. This renders a method that is robust and achieves reproducible results across many months and multiple labs. By conducting several proof-of-concept experiments, we were able to demonstrate the versatility of the cDICE method: from reconstitution of an actin cortex, to encapsulation of a cell-free expression system, membrane-anchored DNA nanostructures, and entire *E. coli* bacteria, these experiments open up a portal to generating GUVs with contents of ever-greater complexity. In the future, additional modifications by changing experimental parameters such as capillary size, rotation speed, chamber design, *etc.* can be made to further extend the possibilities of cDICE and perform experiment-specific optimization. This way, cDICE displays promising potential to become a standard method for the synthetic biology, biochemistry and biophysics communities in the future.

## Methods

### Design and fabrication of the spinning device/rotational chambers

The cDICE device was designed and developed in-house at AMOLF. A 15-Watt Maxon EC32 motor (part number 353400) served as the rotating component of the apparatus, providing a wide range of rotation speeds (from 200 rpm up to 6000 rpm) and allowing precise speed ramps for controlled speeding up and slowing down of rotation. This is especially important to avoid mixing of the solutions after experiments, which would lead to lipid debris in the outer aqueous solution, and to avoid disruption of the formed GUVs. Translucent, cylindrical chambers were designed and printed in-house (Stratasys Objet260 Connex3; Veroclear™ printing material). The chambers measure 38 mm in diameter, have an inner height of 7.4 mm, and include a circular opening of 15 mm in diameter in the top to allow facile access to the solutions with the capillary. The respective designs for rotation chambers and cDICE device are available on GitHub (https://github.com/GanzingerLab). The other two labs at TU Delft used similar devices.

### General cDICE experimental workflow

Synthetic fused silica capillary tubing (TSP 100/050/025 375, Molex) was employed due to its highly smooth inner surface, allowing a controlled flow of inner aqueous solutions. It was cut to a length of several centimetres and attached to a short piece of flexible microbore tubing (Microbore Tubing, 0.020” × 0.060” OD, Cole-Parmer GmbH) using two-component epoxy glue. Using a hollow piece of metal, the capillary tubing was then bent so it could be inserted horizontally into the rotational chamber. To inject the solutions, this set-up was connected to a 250 µL glass syringe (SGE Gas Tight Syringe, luer lock, Sigma-Aldrich) using a shortened needle as connector (Hamilton Needle, Metal hub, needle size 22 ga. blunt tip, Sigma-Aldrich). PEEK capillary tubing (PEEK tubing, 1/32” OD × 0.10 mm ID, BGB Analytik) was used in experiments when explicitly specified. The encapsulation solutions contained 18.5% v/v OptiPrep™ (density gradient medium with a density of 1.320 g mL^−1^) to increase the density. Unless specified otherwise, the outer aqueous phase was a solution of glucose in MilliQ (concentration adjusted to reach a 10 – 20 mOsm higher osmolarity compared to the inner aqueous solution). In a typical experiment, the encapsulation solution was loaded into the set-up, rotation was started, 700 µL of outer aqueous solutions was inserted, followed by 5.5 mL of the lipid-in-oil dispersion. The capillary was then inserted horizontally in the oil layer, until it was visibly embedded. The solution was injected using a syringe pump at a rate of 25 µL min^−1^, unless specified otherwise. The system was spun for a predetermined time depending on the encapsulation volume. Rotation speed ranged from 1000 rpm to 2700 rpm and the capillary diameter from 25 um to 100 µm depending on the experiment type, with 1900 rpm and 100 µm being considered the default values. After every experiment, the chamber was tilted and excess oil was removed. The GUVs were then allowed to sink to the bottom of the rotation chamber for 10 minutes, after which they were harvested using a cut pipette tip and transferred to an observation chamber. Glass coverslips were passivated using 1 mg mL^−1^ beta-casein in MilliQ water. Room humidity was kept around 30 - 40% using a dehumidifier (TTK 71 E Dehumidifier, Trotec). The other two labs used a similar workflow, based on this main protocol.

### Preparation of lipid-in-oil dispersions

1,2-distearoyl-sn-glycero-3-phosphoethanolamine-N-[methoxy(polyethylene glycol)-2000] (18:0 PEG2000 PE), 1,2-dioleoyl-sn-glycero-3-phosphoethanolamine-N-(lissamine rhodamine B sulfonyl (18:1 Liss Rhod PE), 18:1 1,2-dioleoyl-sn-glycero-3-phophocholine (DOPC), 1,2-dioleoyl-sn-glycero-3-[(N-(5-amino-1-carboxypentyl)iminodiacetic acid)succinyl] (nickel salt) (DGS-NTA(Ni)), and 1,2-dioleoyl-sn-glycer-3-phosphoethanolamine-N-(lissamine rhodamine B sulfonyl) (rhodamine-PE) were purchased from Avanti Polar Lipids. ATTO 488 and ATTO 655 labelled 1,2-dioleoyl-sn-glycero-3-phosphoethanolamine (DOPE) were obtained from ATTO-TEC. Stock solutions in chloroform were stored at - 20 °C. The lipids were mixed in the desired molar ratio in a 20 mL glass screw neck vial to obtain a final concentration of 0.2 mg mL^−1^. After desiccation using a gentle nitrogen flow, the vial was brought inside a glovebox, where the lipid film was resuspended in 415 µL of chloroform (Uvasol®, Sigma-Aldrich) or n-decane (99+%, pure, Acros Organics). A mixture of 5.2 mL silicon oil (viscosity 5 cst (25 °C), Sigma-Aldrich) and 1.3 mL mineral oil (BioReagent, Sigma-Aldrich) was then dropwise added to the lipids while vortexing. For the lipid dispersion in mineral oil, 6.5 mL of mineral oil (BioReagent, Sigma-Aldrich) was used instead. After tightly closing the vial and securing the seal with Parafilm®, the lipid-in-oil dispersion was vortexed an additional 2.5 min and sonicated in a bath sonicator for 15 min while keeping the bath temperature below 40 °C. The mixtures were used the same day in experiments.

### UV-VIS absorbance measurements

Turbidity measurements were performed by UV-VIS absorbance using a Denovix DS-11 spectrophotometer. Lipid-in-oil dispersions were prepared as described above and used directly for absorbance measurements. For each measurement, a cuvette (Ultra-micro UV cuvettes, Brand) was filled with 100 µL of lipid-in-oil dispersion and the absorbance at 350 nm was measured thrice. Prior to each measurement, a blank was taken using the corresponding oil or oil mix.

### Pendant drop measurements

Pendant drop measurements were performed using a DSA 30S drop shape analyser (Kruss, Germany) and analysed with the Kruss Advanced software. For each measurement, a lipid-in-oil dispersion containing 100% DOPC was prepared in an identical manner as for cDICE experiments. Directly after vortexing, the mixture was divided over three glass 1.0 mm cuvettes (Hellma Analytics). In each cuvette, a 30 µL droplet containing G-buffer (5 mM tris(hydroxymethyl)aminomethane hydrochloride (Tris-HCl) pH 7.8 and 0.1 mM calcium chloride (CaCl_2_) and 18.5% v/v OptiPrep™ was formed with a rate of 5 µL s^−1^ using an automated dosing system from a hanging glass syringe with needle diameter of 1.060 mm (Hamilton). Immediately when the droplet reached its final volume, 100 frames of the droplets shape were first acquired at a frame rate of 5 fps after which another 500 frames were taken with 1 fps. The droplet contour was automatically detected and fitted with the Young-Laplace equation to yield the interfacial tension. For measurements in dehumidified conditions, a dehumidifier was switched on at least 1 hour prior to the measurement. The lipid-in-oil dispersion was continuously mixed during each measurement using a magnetic stirrer. In several experiments, interfacial tension decreased very rapidly causing the droplet to detach before the end of the measurement.

### Alpha-hemolysin

DOPC (95.4 mol%), DGS-NTA(Ni) (2.5 mol%), and rhodamine-PE lipids (0.1 mol%) were used for preparation of the lipid-in-oil dispersion as described earlier. GUVs encapsulating F-buffer (20 mM Tris-HCl pH 7.4, 50 mM potassium chloride (KCl), 2 mM magnesium chloride (MgCl_2_), 0.5 mM adenosine triphosphate (ATP) and 1mM dithiothreitol (DTT)), 18.5% v/v OptiPrep™, and 5 µM Alexa Fluor 488 (Thermo Fischer) were produced in a 200 mM glucose solution. After production, 50 µl of GUV solution was collected from the bottom of the rotating chamber and deposited on a custom-built observation chamber. Separately, a buffered solution (80 mM Tris pH 7.4 and 240 mM glucose) was mixed with a 4 mg mL^−1^ 4 kDa polyisocyanide hydrogel solution^40^ in a 1:1 volume ratio, and 50 µl of the resulting solution was quickly added to the GUVs. The hydrogel was used to immobilize the GUVs, facilitating extended time-lapse imaging. After a few minutes, 2 µl of 12 µM alpha-hemolysin solution (100 mM Tris-HCl pH 7.5, 1 M sodium chloride (NaCl), 7.5 mM desthiobiotin (DTB)) was added to the observation chamber. Fluorescence intensity was analysed manually using ImageJ and results plotted with MATLAB.

### G-actin encapsulation

DOPC and ATTO 655 DOPE were mixed in a 99.9:0.1 molar ratio to prepare the lipid-in-oil dispersion. 100 µL of actin (4.4 µM, 9% labelled with Alexa Fluor 488) in G-buffer (5 mM Tris-HCl pH 7.8, 0.1 mM CaCl_2_, 0.02 mM ATP and 4 mM DTT) and 18.5% v/v OptiPrep™ was encapsulated in every experiment, only varying rotation speed and capillary size. For a capillary size of 25 µm, the flow rate was lowered to 2.5 µL min^−1^, to reduce the pressure in the capillary set-up. The encapsulated volume was reduced to 50 µL in these experiments. GUVs were produced in an outer aqueous solution containing approximately 85 mM glucose in MilliQ water. Actin was purchased from Hypermol and Alexa Fluor 488-labelled actin was obtained from Invitrogen. All proteins were handled according to instructions provided by the manufacturer. GUVs were imaged in the outer aqueous solution using confocal microscopy, 50 µL of GUV solution was deposited on a custom-made glass coverslip and covered. Microscopy was performed using a Nikon A1R-MP confocal microscope, using a Plan APO IR 60× water immersion objective. The 561 nm (laser power 1.0) and 488 nm (laser power 1.0) laser lines were used in combination with appropriate emission filters to image the ATTO 655-labelled DOPE membrane and Alexa Fluor 488-labelled G-actin, respectively.

### Data analysis of GUV images

GUV size and inner intensity (Fig. 1d, Fig. 3a, c, and SI Fig. 3) were obtained from Z-stack images that were processed using custom-written Python software. The software performs feature tracking in each frame in three consecutive steps. First, the Canny edge detection algorithm^76^ is applied, then filling of the detected edges is achieved by applying the binary hole filling function from the ndimage module of the SciPy package^77^, and lastly these features in each frame are located using the measure module of the scikit-image package^78^ for Python. The located features are linked together in a final step to group points belonging to the same GUV along the frame-axis. The radius of the GUVs was determined from the frame where the detected feature was largest and the inner intensity was also obtained from that respective frame and feature. User-based filtering was applied afterwards to discard multilamellar structures, aggregates or similar. The software is available on GitHub (https://github.com/GanzingerLab). The intensity was normalized to the mean of the distribution.

### PURE system encapsulation

The codon-optimized construct encoding for *meYFPco-LL-spinach* (enhanced yellow fluorescent protein) described in Van Nies *et al.*^79^ was used. The sequence is codon-optimized for expression in the PURE system, and the template includes the T7 promoter and terminator. A linear DNA template was employed to observe fluorescence readout of the level of synthesized protein. The linear DNA construct was obtained by polymerase chain reaction (forward primer: GCGAAATTAATACGACTCACTATAGGGAGACC, reverse primer: AAAAAACCCCTCAAGACCCGTTTAGAGG). Amplification products were checked on a 1% agarose gel and were purified using the Wizard PCR clean-up kit (Promega). DNA concentration and purity were measured using a ND-1000 UV-Vis Spectrophotometer (Nanodrop Technologies).

The full sequence of the *meYFPco-LL-spinach* linear construct is: 5’GCGAAATTAATACGACTCACTATAGGGAGACCACAACGGTTTCCCTCTAGAAATAATTTTGTTTAACTTTAAGAAGGAGATATACATATGCGGGGTTCTCATCATCATCATCATCATGGTATGGCTAGCATGACTGGTGGACAGCAAATGGGTCGGGATCTGTACGAC GATGACGATAAGGATCCGATGGTTAGCAAAGGCGAAGAACTGTTTACGGGCGTGGTGCCGATTCTGGTGGAACTGGACGGCGACGTGAACGGTCACAAATTCAGCGTTTCGGGCGAAGGTGAAGGCGATGCGACCTATGGTAAACTGACGCTGAAATTTATTTGCACCACCGGTAAACTGCCGGTGCCGTGGCCGACCCTGGTTACCACGTTTGGTTATGGCCTGCAGTGTTTCGCGCGCTACCCGGATCATATGAAACAACACGACTTTTTCAAATCTGCCATGCCGGAAGGTTATGTGCAGGAACGTACGATTTTCTTTAAAGATGACGGCAACTACAAAACCCGCGCAGAAGTCAAATTTGAAGGTGATACGCTGGTGAACCGTATTGAACTGAAAGGCATCGATTTCAAAGAAGACGGTAATATCCTGGGCCATAAACTGGAATACAACTACAACTCCCACAACGTTTACATCATGGCAGATAAACAGAAAAACGGTATCAAAGTCAACTTCAAAATCCGCCATAACATCGAAGATGGCTCAGTGCAACTGGCTGACCACTACCAGCAAAACACCCCGATCGGTGATGGCCCGGTTCTGCTGCCGGACAATCATTATCTGAGCTACCAGTCTAAACTGAGTAAAGATCCGAACGAAAAACGTGACCACATGGTCCTGCTGGAATTTGTGACGGCGGCTGGTATTACGCTGGGCATGGATGAACTGTATAAATGAAAGCTTCCCGGGAAAGTATATATGAGTAAAGATATCGACGCAACTGAATGAAATGGTGAAGGACGGGTCCAGGTGTGGCTGCTTCGGCAGTGCAGCTTGTTGAGTAGAGTGTGAGCTCCGTAACTAGTCGCGTCGATATCCCCGGGCTAGCATAACCCCTTGGGGCCTCTAAACGGGTCTTGAGGGGTTTTTT-3’.

DOPC and rhodamine-PE were used in a 99.9:0.1 molar ratio for the lipid-in-oil dispersion, 0.01 mol% of 18:0 PEG2000 PE was used when explicitly mentioned. PURE*frex*2.0 (GeneFrontier Corporation, Japan) was utilized following storage and handling instructions provided by the supplier. Linear DNA template was added at a concentration of 5 nM. Reactions of 40 µL were assembled in test tubes and supplemented with 5% v/v OptiPrep™ (higher ratios negatively interfered with the PURE reaction) and kept on ice. GUVs were produced in an outer aqueous solution composed of 220 mM glucose in MilliQ water. The flow rate was kept at 2.5 µl min^−1^ for 8 minutes in total, given the limited availability of inner aqueous solution. After production, 25 µl of GUV solution was transferred to the observation chamber, together with 25 µl of additional outer aqueous solution composed of 35 mM glucose and 50% v/v PURE buffer. YFP expression was monitored at 37 °C by confocal imaging using a Nikon A1R Laser scanning confocal microscope equipped with an SR Apo TIRF 100× oil-immersion objective. The 561 nm (laser power 5.0) and 488 nm (laser power 20.0) laser lines were used in combination with appropriate emission filters to image the rhodamine-PE membrane and YFP, respectively. The software NIS (Nikon) was used for image acquisition and the settings were maintained for all experiments. Samples were mounted on a temperature-controlled stage maintained at 37 °C during imaging up to five hours.

Image analysis was carried out in MATLAB version R2020b using the script published in Blanken *et al*.^80^. Briefly, the script reads the split-channel tiff files, identifies the GUVs, indexes them, and then stores the indexed variables in the data file. The script uses a sharpening filter on the rhodamine-PE image, the GUV lumen is determined by a flood filling step followed by a binarization phase with a cut-off of 200. An erosion step was conducted to filter segments relative to lipid aggregates and other sources of noise. Any segments with a circularity of less than 0.5 or greater than 2 have been excluded. For each GUV, average rhodamine-PE intensity, average YFP intensity and YFP intensity variance were determined. The box plots of the YFP intensity in the lumen were also generated in MATLAB version R2020b.

### Actin cortex

GUVs were prepared using a mixture of DOPC and DGS-NTA(Ni) lipids in a 50:1 molar ratio. Actin (4.4 µM, 9% labelled with Alexa Fluor 647), profilin (3.3 µM), Arp2/3 (100 nM), and VCA (0.6 µM) were added to a solution containing F-buffer (20 mM Tris-HCl pH 7.4, 50 mM KCl, 2 mM MgCl_2_, 0.5 mM ATP and 1 mM DTT) and 18.5% v/v OptiPrep™. To minimize photobleaching, an oxygen-scavenger system^81^ (1 mM protocatechuic acid (PCA) and 50 nM protocatechuate-3,4-dioxygenase (PCD)) was also added to the solution. GUVs were produced in an outer aqueous solution containing 200 mM glucose in MilliQ water. After production, 25 µl of GUV solution was collected from the bottom of the rotating chamber and deposited on a custom-built observation chamber, to which an additional 25 µl of a buffered solution (40 mM Tris pH 7.4 and 125 mM glucose) was added. Unless specified otherwise, all chemicals were purchased from Sigma-Aldrich. All proteins, except VCA, which was purified in-house, were purchased from Hypermol and dissolved according to instructions provided by the manufacturer. G-actin was dialyzed in G-buffer (5 mM Tris-HCl pH 7.8 and 0.1 mM CaCl_2_) before storage at - 80 °C.

### Alpha-hemolysin

DOPC (95.4 mol%), DGS-NTA(Ni) (2.5 mol%), and rhodamine-PE lipids (0.1 mol%) were used for preparation of the lipid-in-oil dispersion as described earlier. GUVs encapsulating F-buffer (20 mM Tris-HCl pH 7.4, 50 mM potassium chloride (KCl), 2 mM magnesium chloride (MgCl_2_), 0.5 mM ATP and 1mM DTT), 18.5 % v/v OptiPrep™, and 5 µM Alexa Fluor 488 (Thermo Fischer) were produced in a 200 mM glucose solution. After production, 50 µl of GUV solution was collected from the bottom of the rotating chamber and deposited on a custom-built observation chamber. Separately, a buffered solution (80 mM Tris pH 7.4 and 240 mM glucose) was mixed with a 4 mg mL^−1^ 4 kDa polyisocyanide hydrogel solution^40^ in a 1:1 volume ratio, and 50 µl of the resulting solution was quickly added to the GUVs. The hydrogel was used to immobilize the GUVs, facilitating extended time-lapse imaging. After a few minutes, 2 µl of 12 µM alpha-hemolysin solution (100 mM Tris-HCl pH 7.5, 1 M sodium chloride (NaCl), 7.5 mM desthiobiotin (DTB)) was added to the observation chamber.

### DNA origami nanostructures encapsulation

The DNA origami design was adapted from Jeon *et al.*^62^ by removing the 3’ sequence (“sticky ends”) mediating multimerization, thus keeping them monomeric. An additional 12 nt sequence was added at the 5’ end to allow binding to the membrane via a cholesterol-oligonucleotide anchor. Nanostructures were folded by thermal annealing (from 95 °C to 23 °C, - 0.5 °C min^−1^) and used at 1 µM in buffered solution (50 mM Tris pH 7.0, 2 mM MgCl_2_ and 200 mM sucrose). Right before encapsulation, 2 µM of cholesterol-oligonucleotides were added to this buffer. As an outer aqueous phase, 50 mM Tris pH 7.0, 2 mM MgCl_2_ and 200 mM glucose was used. Experiments were performed using PEEK capillary tubing.

### SUV encapsulation

SUVs were prepared using DOPC and ATTO 488 DOPE in a 99:1 molar ratio. Under gentle nitrogen flow, chloroform was evaporated to obtain a homogenous lipid film. The lipid film was then desiccated for a minimum of three hours to remove any remaining solvent traces, after which it was rehydrated in phosphate-buffered saline buffer (PBS buffer) at 4 mg mL^−1^ by vortexing. Afterwards the solution was sonicated in aliquots of 20 µL for 2x 30 min. It was then diluted to 0.5 mg mL^−1^ for further use. DOPC and ATTO 655 DOPE were used in a 99.9:1 molar ratio for the lipid-in-oil dispersion. For encapsulation, the SUVs were diluted 10× in PBS buffer and 18.5% v/v OptiPrep™ was added. The outer aqueous phase consisted of 313 mM glucose in MilliQ water.

### Bacteria encapsulation

DOPC, 18:1 Liss Rhod PE, and 18:0 PEG2000 PE were used in a 98.9:0.1:1 molar ratio for the lipid-in-oil dispersion. A saturated lysogeny broth (LB) culture of *Escherichia coli* expressing green fluorescent protein (GFP-HU) was centrifuged and the pellet resuspended in a buffered solution (50 mM Tris pH 7.5, 5 mM NaCl and 200 mM sucrose) and used for encapsulation. As an outer aqueous phase, 50 mM Tris pH 7.5, 5 mM NaCl and 200 mM glucose was used. cDICE experiments were performed using PEEK capillary tubing.

## Acknowledgements

We thank Paul Kouwer (Radboud University) for the kind gift of the polyisocyanide gel, and we thank Josef Melcr and Siewert-Jan Marrink for useful discussions. We acknowledge the financial support by the “BaSyC - Building a Synthetic Cell” Gravitation grant (024.003.019) of the Netherlands Ministry of Education, Culture and Science (OCW) and the Netherlands Organization for Scientific Research (G.H.K., C. Dekker, and C. Danelon) and NWO-WISE funding (K.A.G.).

## Author contributions

K.A.G. and G.H.K. designed and directed the project; L.V.d.C., F.F., L.v.B., N.D.F, E.G. and S.B. performed the experiments; L.V.d.C., F.F. and L.v.B. drafted the manuscript and designed the figures; all authors provided critical feedback and helped shape the research and manuscript.

---

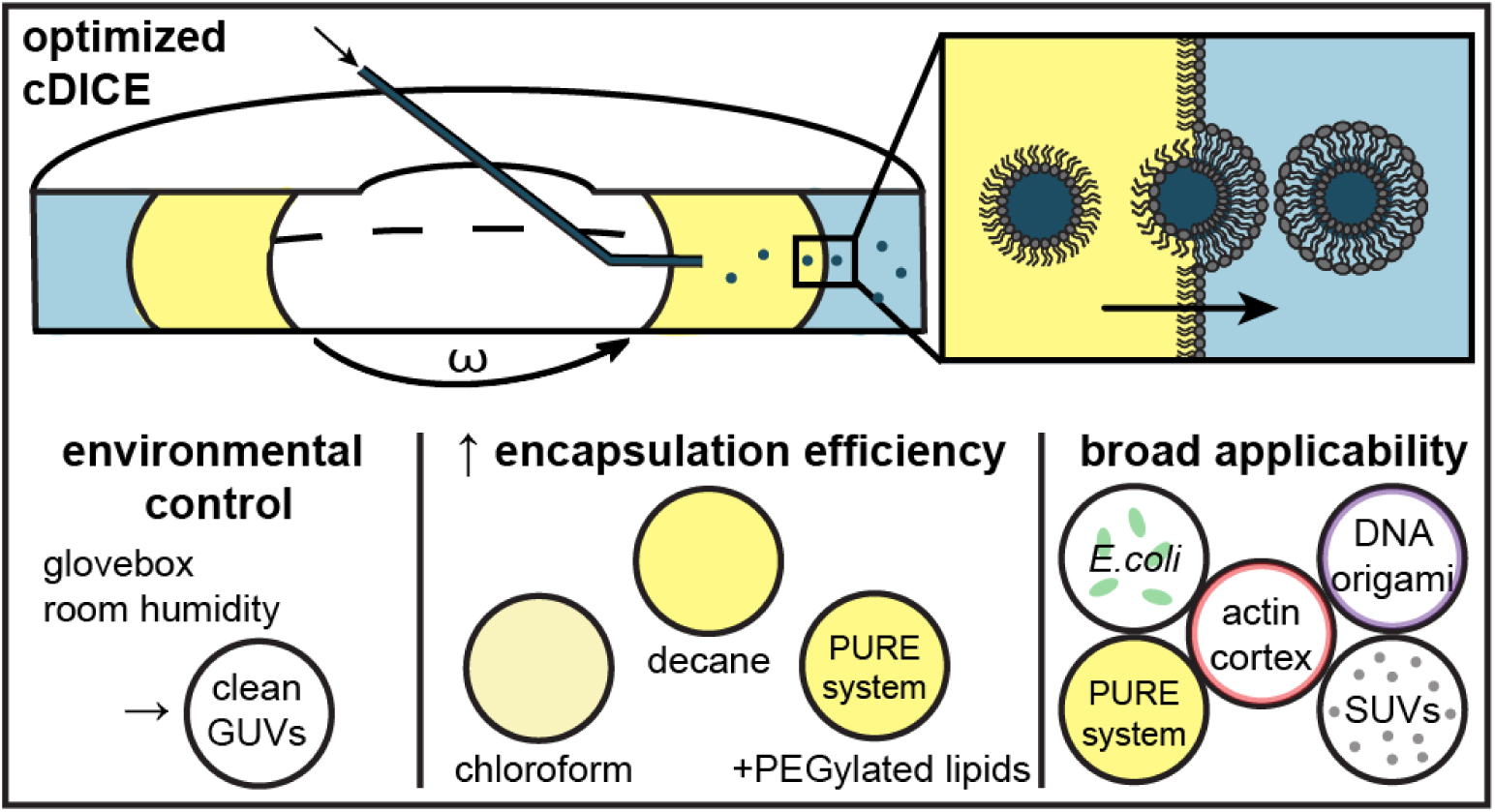
For Table of Contents Only.

## Notes

### Competing Interest Statement

The authors have declared no competing interest.

