## Supplementary Information for "Optimized cDICE for efficient reconstitution of biological systems in giant unilamellar vesicles"

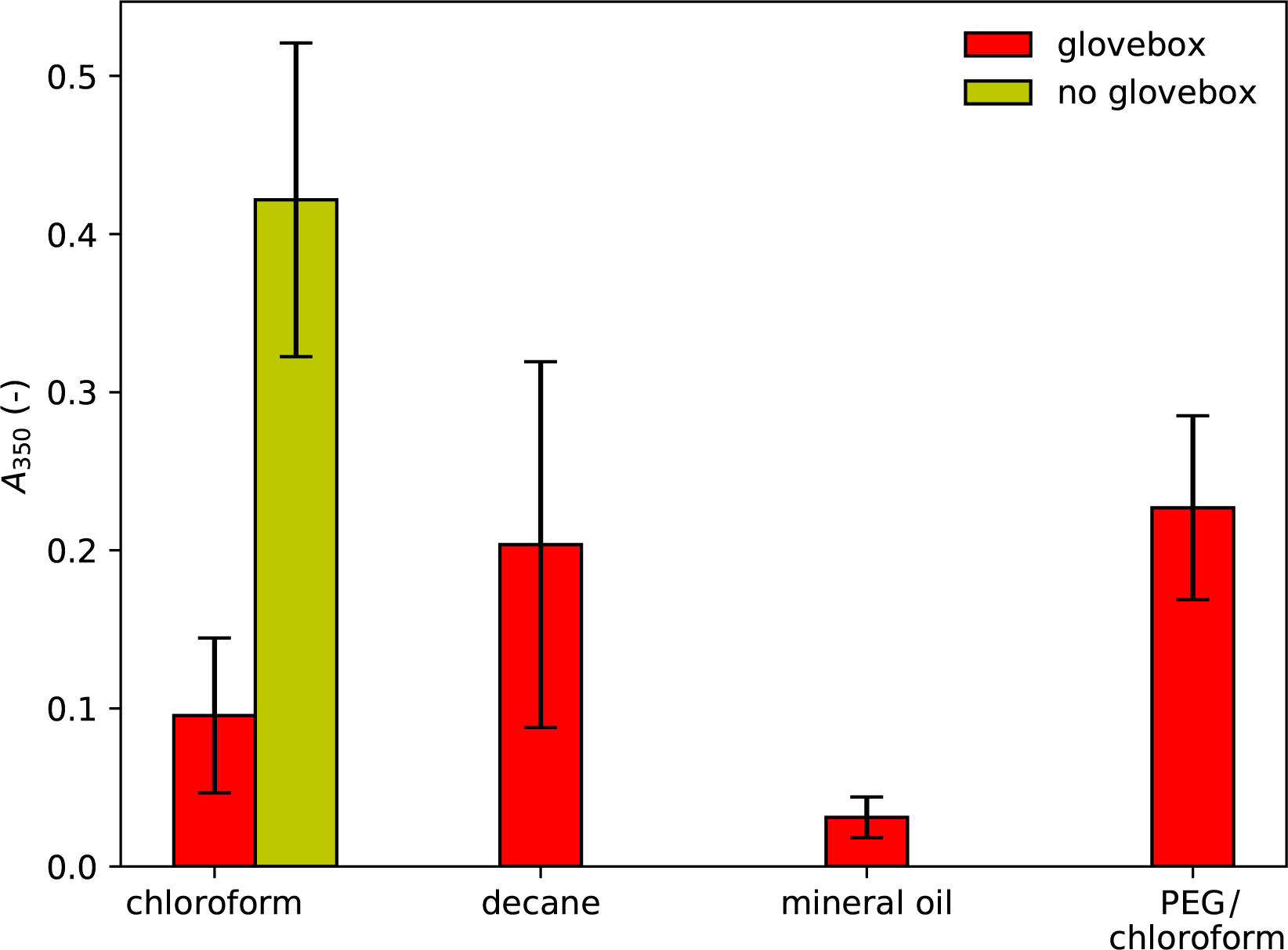

**SI Figure 1.** UV-VIS absorbance of different lipid-in-oil mixtures.

Absorbance at 350 nm of lipid-in-oil dispersions measured right after preparation. The different lipid-in-oil mixtures consist of: dispersed lipid aggregates using chloroform or decane in silicone oil:mineral oil 80:20, a lipid-chloroform solution in mineral oil only, and a chloroform-based lipid-in-oil dispersion with 0.01 mol% of PEGylated lipids. Red bars indicate dispersions that were prepared in the glovebox. The green bar shows the turbidity of the chloroform-based dispersion prepared outside of the glovebox. Data represents three measurements on at least two individual preparations with standard deviation.

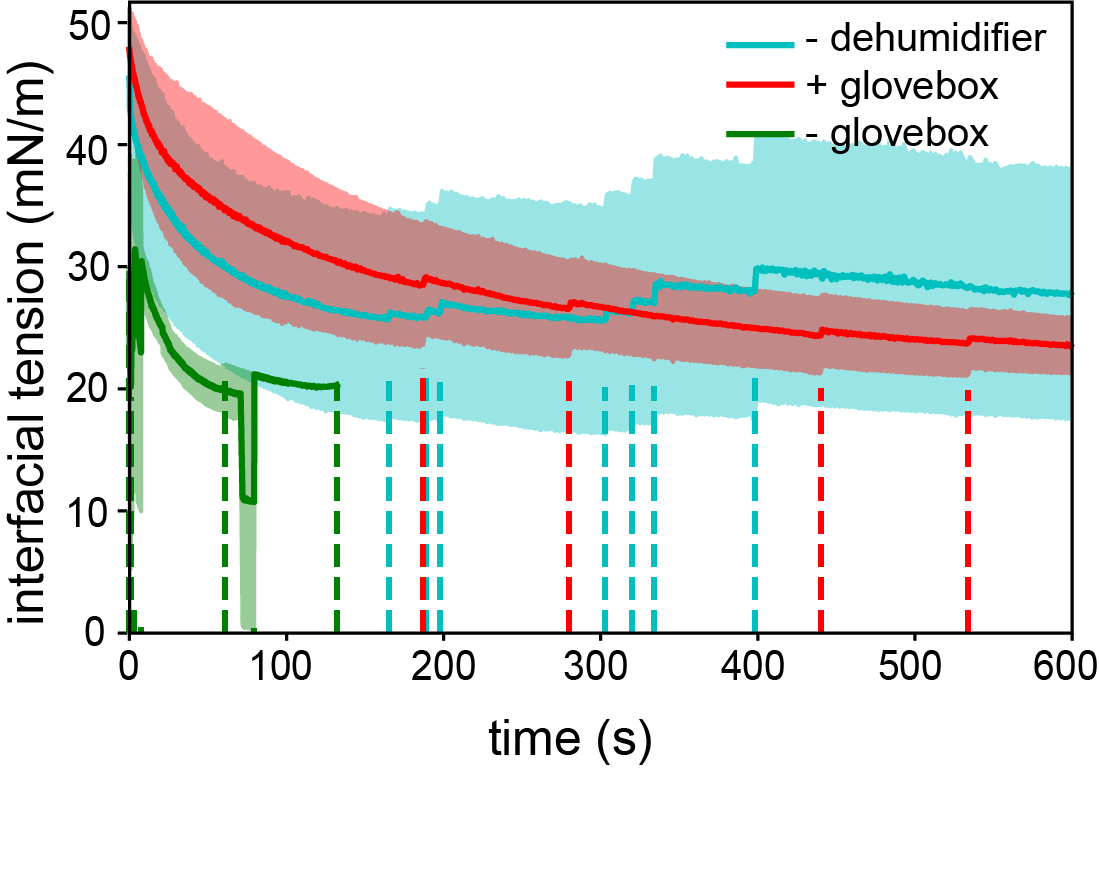

**SI Figure 2.** Influence of environmental conditions on lipid adsorption kinetics.

Interfacial tension decrease measured for a pendant droplet of G-buffer in different lipid-in-oil mixtures. Solid lines show averaged data with standard deviation for a chloroform-based lipid-in-oil dispersion prepared inside a glovebox with dehumidifier in the experiment room (red, n = 13), outside a glovebox with dehumidifier in the experiment room (green, n = 9), and inside a glovebox without dehumidifier in the experiment room (blue, n = 11). The dashed lines indicate individual events where the droplet fell off, which gave rise to apparent jumps in the averaged curves.

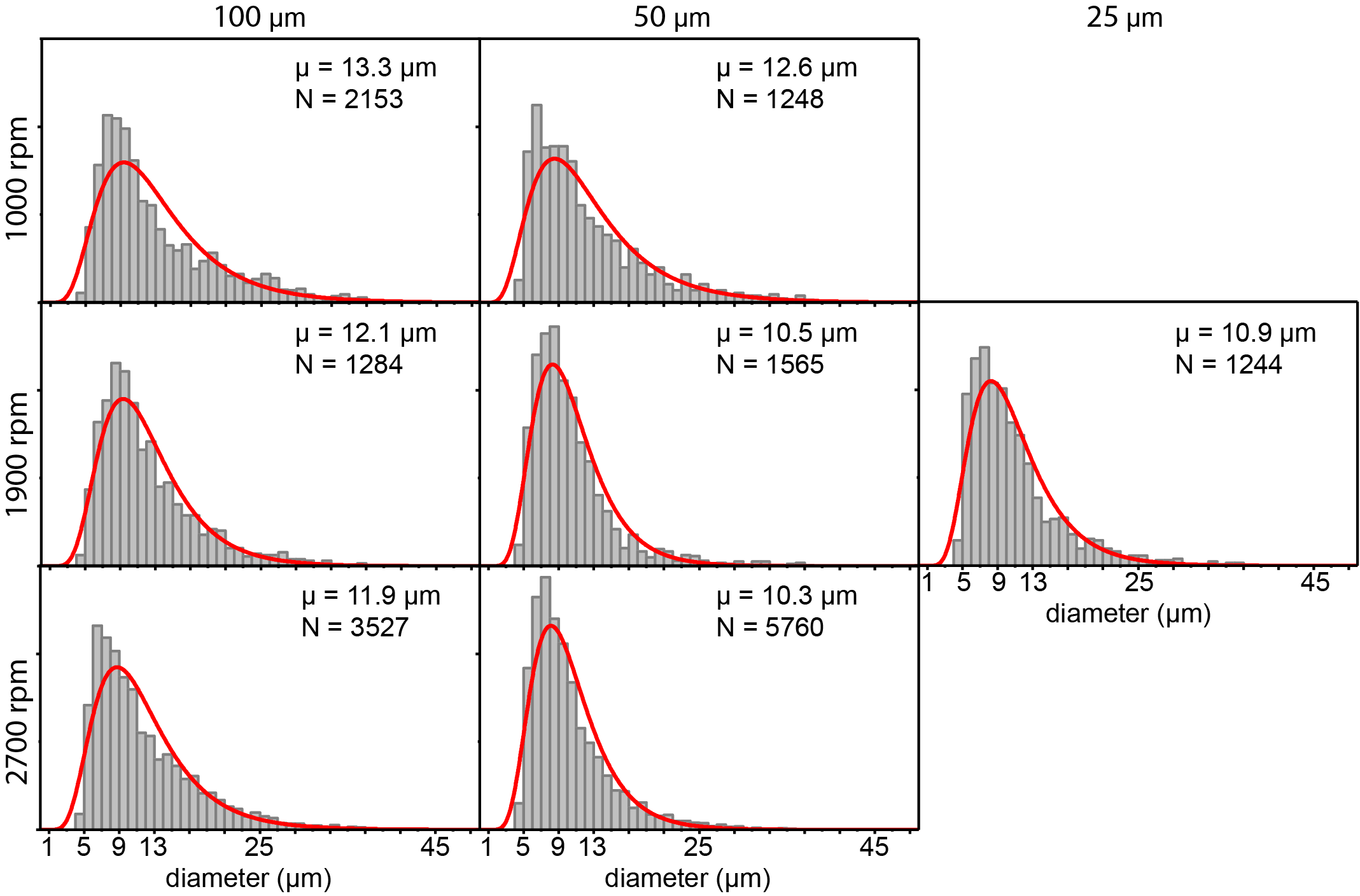

**SI Figure 3.** Size distributions for different capillary sizes and rotation speeds.

Size distribution of GUVs made of DOPC lipids, using capillary sizes 25 µm, 50 µm, and 100 µm, and rotation speeds 1000 rpm, 1900 rpm, and 2700 rpm. The individual graphs represent pooled data for three experiments. The distributions are fitted to a log-normal function (red curves).

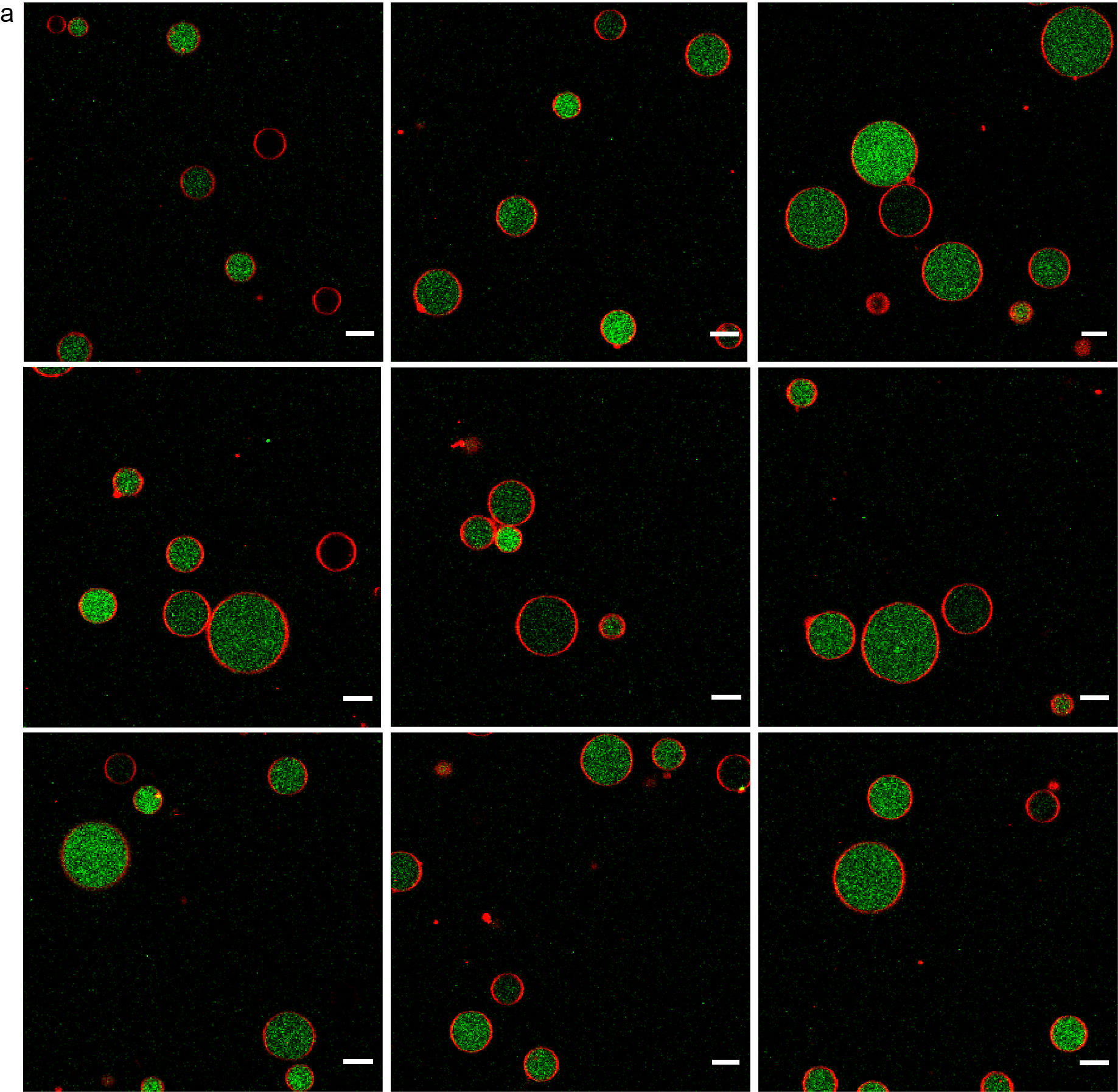

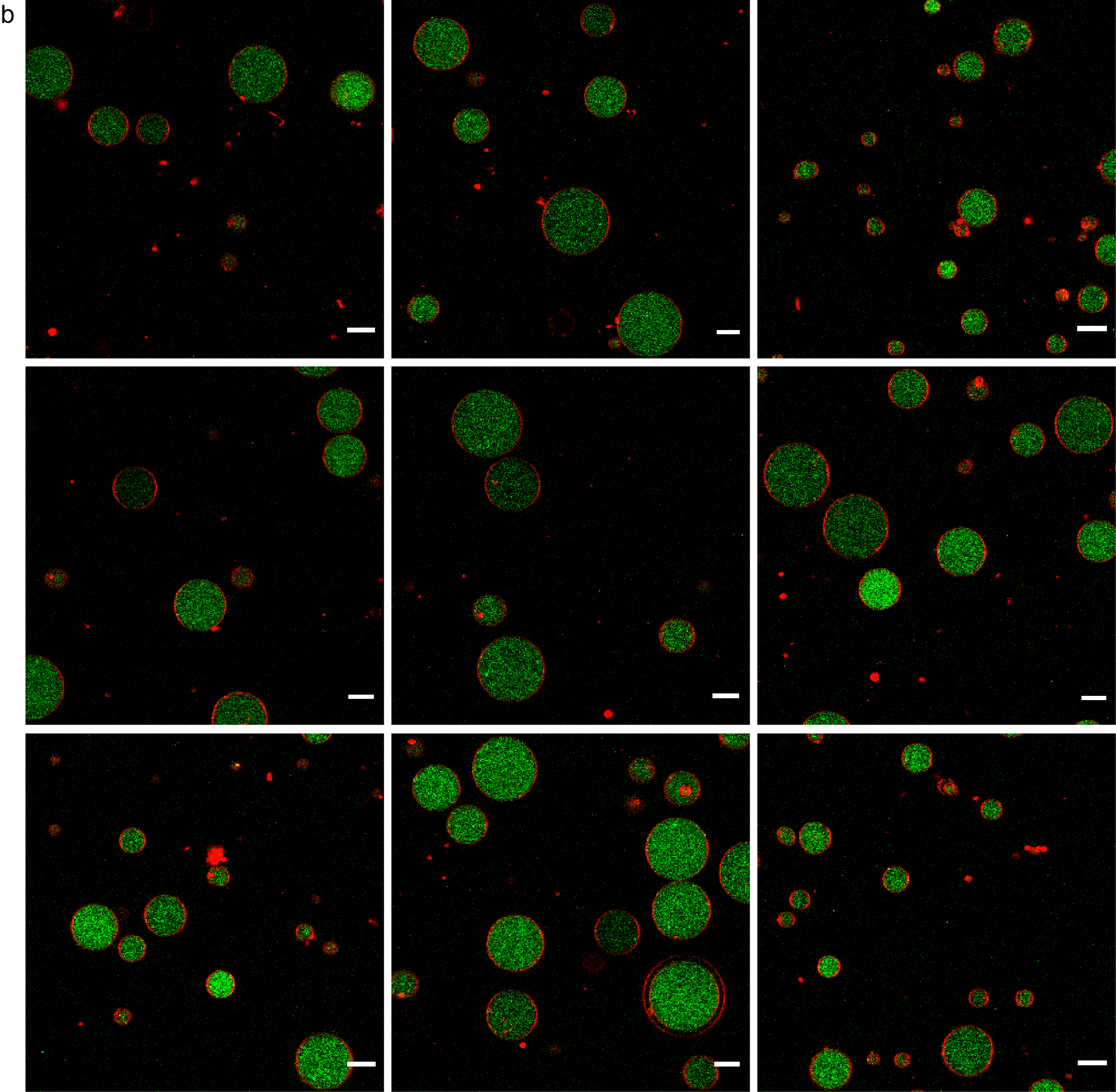

**SI Figure 4.** Representative fields of view of GUVs encapsulating G-actin using:

a. Chloroform-based lipid-in-oil dispersion
b. Decane-based lipid-in-oil dispersion

Scale bars indicate 10 µm. Images were acquired with identical imaging settings.

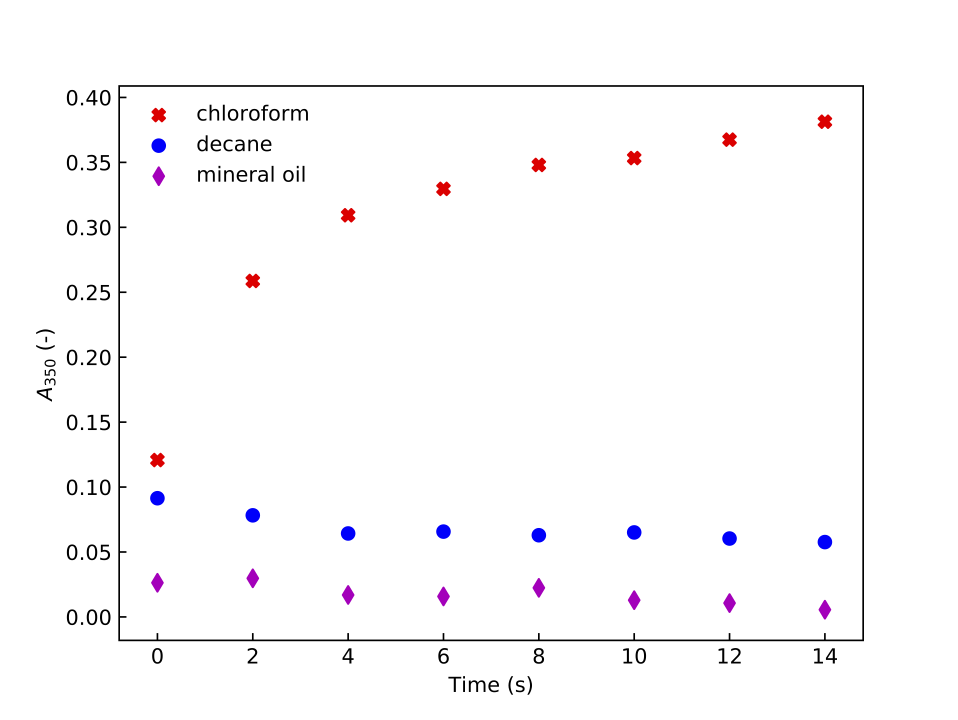

**SI Figure 5.** Time traces of UV-VIS absorbance of different lipid-in-oil mixtures

Absorbance at 350 nm of the lipid-in-oil dispersion was measured over ten minutes after opening of the vial. Measured samples include dispersed lipid aggregates using chloroform or decane in silicone oil:mineral oil 80:20 and a lipid-chloroform solution in mineral oil only. All samples were prepared in the glovebox.

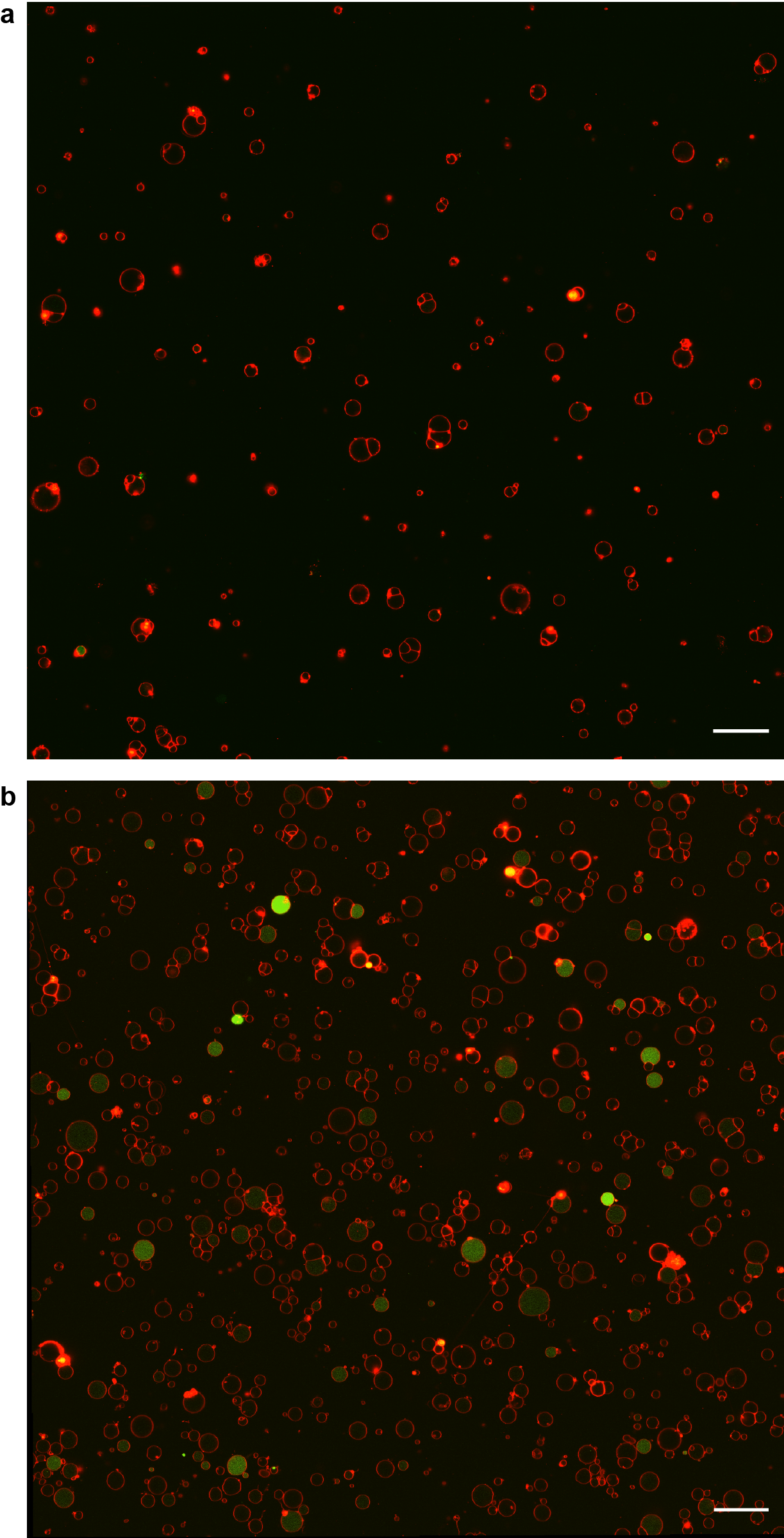

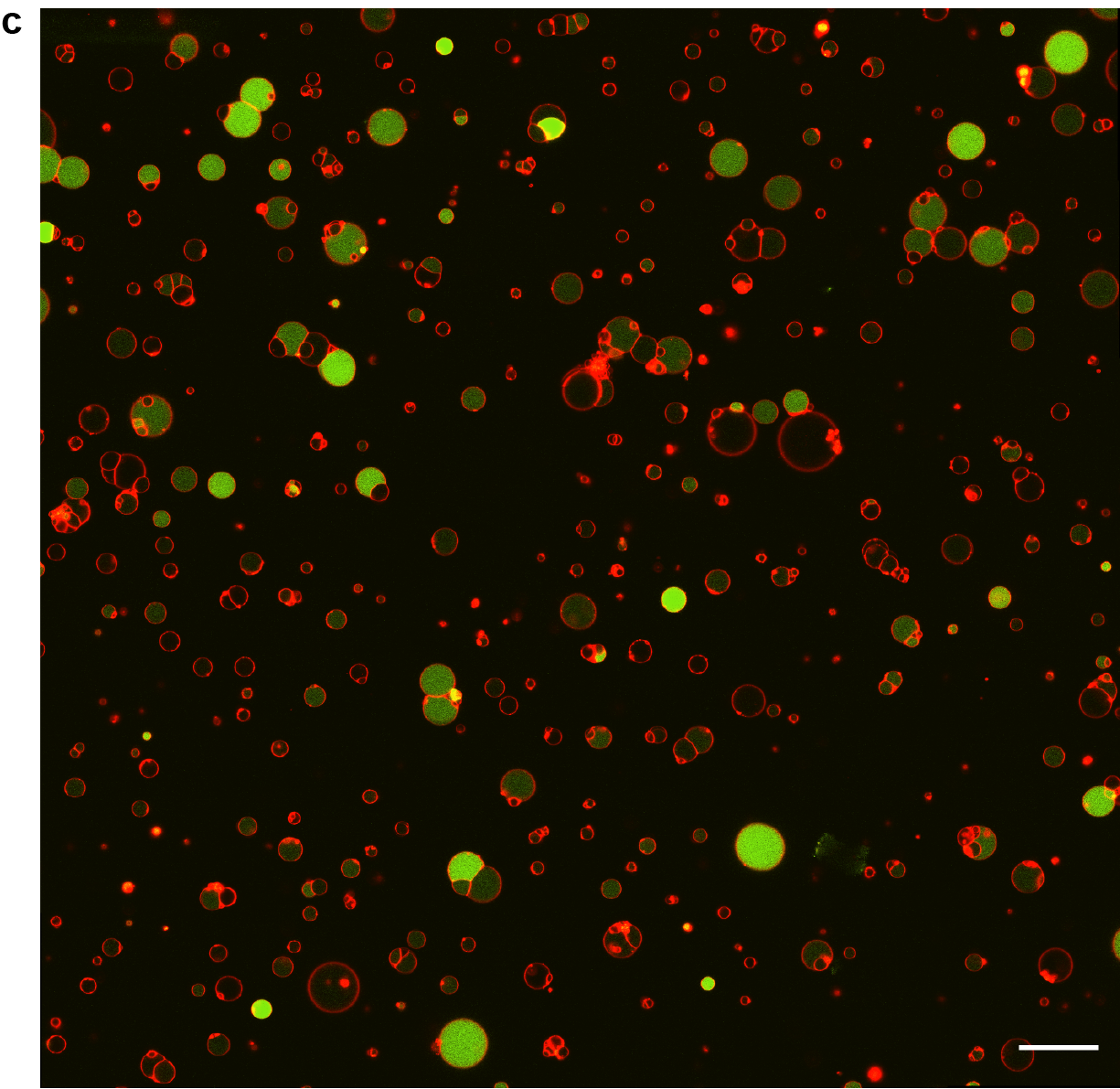

**SI Figure 6.** Representative fields of view of GUVs encapsulating the PURE system.

a. Encapsulation of PURE*frex*2.0 and DNA encoding for YFP using a chloroform-based lipid-in-oil dispersion.
b. Encapsulation of PURE*frex*2.0 and DNA encoding for YFP using a decane-based lipid-in-oil dispersion.
c. Encapsulation of PURE*frex*2.0 and DNA encoding for YFP using a chloroform-based lipid-in-oil dispersion and 0.01 mol% PEGylated lipids.

All pictures have the same size (scale bars indicate 50 µm) and were acquired with	identical imaging settings.

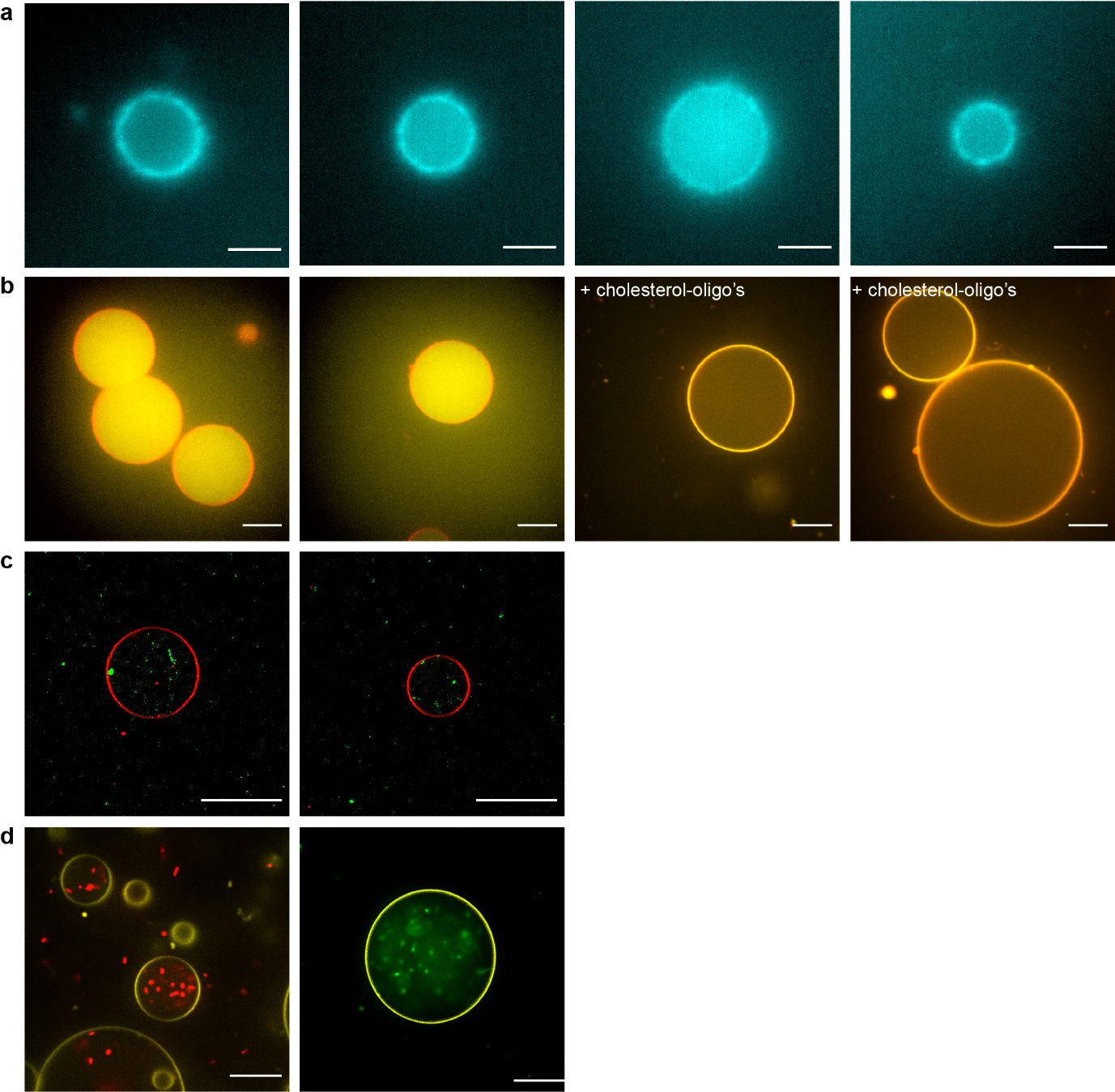

**SI Figure 7.** Representative fields of view of proof-of-concept experiments.

a. Reconstitution of a minimal actin cortex inside a GUV, nucleated at the vesicular membrane by the Arp2/3 complex, the C-terminal VCA domain of WASp, and profilin. Scale bar indicates 5 µm.
b. Encapsulation of DNA origami nanostructures, freely diffusing inside the GUV lumen and capable of membrane localization upon addition of 2 µM of cholesterol-oligonucleotides. Scale bar indicates 15 µm.
c. Encapsulation of SUVs inside GUVs to form a multicompartmentalized system. Scale bars indicate 20 µm.
d. Encapsulation of GFP-HU expressing *E. coli* bacteria. A large number of bacteria could be observed inside the GUV lumen, clearly viable as evident from their motility. Scale bar indicates 20 µm.

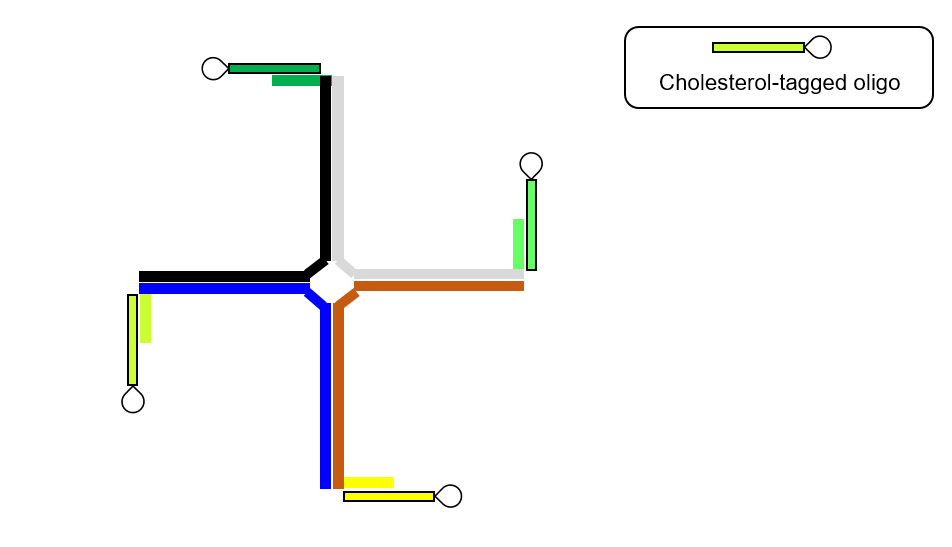

**SI Figure 8.** Schematic of a single DNA nanostructure.

The DNA nanostructure consists of four single-stranded DNA oligonucleotides (grey, black, blue, and brown) that hybridize forming a cross-like shape. They all have a complementary hybridization sequence for the cholesterol-oligonucleotide at their 5’ end.

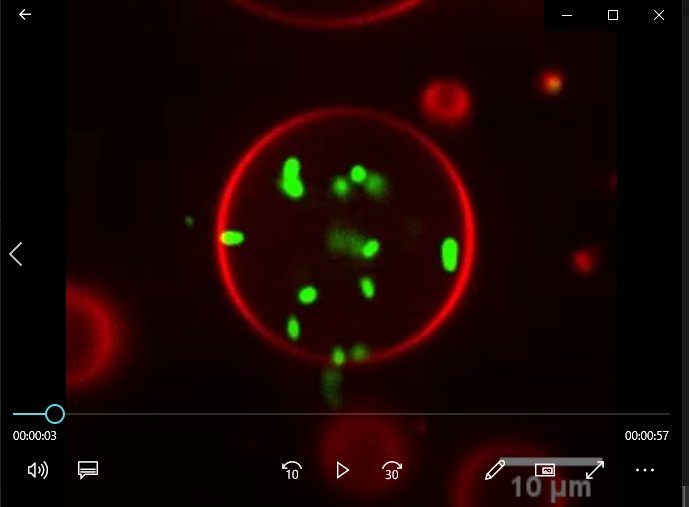

*screenshot*

**SI Movie 1.** Time-lapse movie of GFP-HU-expressing *E. coli* bacteria encapsulated in a GUV.

The GFP-HU-expressing *E. coli* bacteria (green) are clearly mobile inside the GUV (red), indicating their viability. The time per frame is 1 sec, and the movie is displayed in real time.
